# Neuronal expression of E2F4DN restores adult neurogenesis in homozygous 5xFAD mice via TrkB signaling

**DOI:** 10.1101/2024.12.17.628978

**Authors:** Anna Lozano-Ureña, Aina Maria Llabrés-Mas, Morgan Ramón-Landreau, Alberto Fraj-Cebrián, Cristina Sánchez-Puelles, Alberto Garrido-García, Vanesa Cano-Daganzo, Lorena Valdés-Lora, José María Frade

**Author notes:** Correspondence: José María Frade.

## Abstract

**Background:** Alzheimer’s disease (AD) is increasingly recognized as a disorder that disrupts brain plasticity long before extensive neuronal loss occurs, including early impairment of adult neurogenesis. However, the molecular mechanisms underlying these alterations remain poorly understood.

**Methods:** Neuronal expression of E2F4 carrying the Thr249Ala/Thr251Ala mutation (E2F4DN) was achieved using an AAV.PHP.eB vector in the 5xFAD mouse model of AD. Neurogenic potential was assessed by immunohistochemistry and quantitative PCR (qPCR) targeting neurogenesis-associated markers. For *in vitro* experiments, neural stem cells (NSCs) isolated from the subventricular zone (SVZ) were exposed to conditioned media from primary cortical neurons expressing E2F4DN, with or without pharmacological inhibitors. NSC proliferation and differentiation were evaluated by immunocytochemistry and qPCR.

**Results:** Neuronal expression of E2F4DN restored the proliferative and differentiation capacity of NSCs in 5xFAD mice both in vivo and in vitro. Pharmacological inhibition of TrkB signaling abolished the pro-neurogenic effects of E2F4DN-conditioned medium, whereas direct activation of TrkB in NSCs recapitulated these effects.

**Conclusions:** These findings identify a mechanism by which neuronal dysfunction in AD compromises neurogenesis through impaired TrkB-mediated signaling. The results uncover a previously underappreciated paracrine communication between neurons and NSCs and highlight neuronal E2F4DN expression and TrkB modulation as promising therapeutic strategies to counteract early neurogenic deficits in AD.

## Background

Alzheimer’s disease (AD) is a neurodegenerative disorder and the most common cause of dementia worldwide. AD is a multifactorial condition, with multiple etiopathological mechanisms proposed to contribute to its onset and progression. Among these, altered neurogenesis in the two principal neurogenic niches, the subventricular zone (SVZ) lining the lateral ventricles and the subgranular zone (SGZ) of the hippocampal dentate gyrus, have been identified as an early event in disease pathogenesis (1). Hippocampus is a crucial region for learning and memory, and is one of the main structures affected in AD (2). Thus, adult hippocampal neurogenesis has attracted the major focus on AD research; however, alterations in the adult SVZ neurogenesis (ASN) appear presymptomatically (1), being olfactory dysfunction one of the earliest clinical symptoms of AD (3, 4). Consistent with these early changes, olfactory dysfunction has been proposed as a potential tool for early diagnosis (4). Nevertheless, the mechanisms underlying ASN dysfunction in AD and its therapeutic relevance remain incompletely understood. The neurogenic niches contain a cell population, known as neural stem cells (NSCs), responsible to generate new neurons throughout life of the individual. Due to the radial-glia origin of the adult NSCs, these cells share features with astrocytic population such as the expression of the glial fibrillary acidic protein (GFAP) or the glutamate transporter (GLAST). NSCs also express the transcription factor SRY-related HMG box 2 (Sox2) and the neural progenitor markers Nestin and Ascl1 (5). NSCs may exist in a dormant or primed quiescence state, during which they do not divide. Alternatively, they can be activated, leading to rapid cell divisions, although the return to a quiescence state is also possible (6). NSCs in the SVZ, when activated, generate new cells through transit-amplifying progenitors (TAPs or C cells) that rapidly divide into DCX positive neuroblasts (A cells), that migrate through the rostral migratory stream (RMS) to reach the olfactory bulb (OB) where they differentiate into mature interneurons (7). NSCs from the SVZ also generate astrocytes and oligodendrocytes that integrate into the corpus callosum (CC) (8) and striatum (9).

ASN is modulated by several intrinsic and extrinsic factors, including methylation and transcription factors as well as growth factors and neurotransmitters, respectively (1). E2F4 is a transcription factor that plays relevant roles in cell and tissue homeostasis and regeneration, and controls the gene networks affected in AD (10). E2F4 can be phosphorylated by p38^MAPK^ in the conserved Thr248/Thr250 motif (Thr249/Thr251 in mouse E2F4) (11), and this phosphorylation may prevent the homeostatic capacity of E2F4 (10, 12). We have previously proved in transgenic mice that the expression in neurons of a dominant negative form of mouse E2F4 (mE2F4DN) containing Thr249Ala/Thr251Ala mutations prevents the pathological phenotype of 5xFAD mice (13), an Alzheimer mouse model that recapitulates many of the features of human AD (14). The cerebral cortex of 5xFAD transgenic mice carrying a neuron-specific *mE2F4DN* knock-in allele exhibited a transcriptional program consistent with attenuation of the immune response and maintenance of brain homeostasis. This correlated with reduced microgliosis and astrogliosis, modulation of amyloid-β (Aβ) peptide proteostasis, blocking of neuronal tetraploidization, prevention of cognitive impairment and of body weight loss, a known somatic alteration associated with AD (13). In homozygous 5xFAD (h5xFAD) mice, neuron-specific expression by gene therapy of human E2F4DN (Thr248Ala/Thr250Ala) reduced Aβ burden, dampened astrocytosis and microgliosis, abolished neuronal tetraploidization, and prevented cognitive deficits (15). This treatment also reversed other alterations observed in h5xFAD mice such as paw-clasping behavior and body weight loss (15). These effects are consistent with the phosphorylation of E2F4 at Thr249, detected in the brain of h5xFAD mice (15), or the presence of E2F4 in cortical neurons from Alzheimer patients in association with Thr-specific phosphorylation (13); and the finding that both p38^MAPK^ activation and E2F4 phosphorylation are molecular changes found in neurodegenerative diseases (16–18).

TrkB signaling has been postulated as an important factor in the AD pathogenesis and one of the most important endogenous neuroprotective systems (19, 20). Accordingly, the use of TrkB agonists, such as 7,8-dihydroxyflavone (7,8-DHF) or R13, enhanced learning and memory (20–22) and neurogenesis (23) in rodents. More recently, a small TAT-fused peptide has been developed to prevent TrkB cleavage, a phenomenon that occurs more frequently in the AD context (24), rescuing synaptic deficits and cognitive performance in 5xFAD animals (25). We have shown that TrkB signaling is able to modulate ASN (26), while its ligand brain-derived neurotrophic factor (BDNF) has been shown to be expressed in the SVZ neurogenic niche (27). TrkB can also transduce signals from NT4/5 (28), another member of the neurotrophin family that has been shown to be expressed by NSCs (29). Therefore, NT4/5 might participate in ASN. However, little is known about the role of NT4/5 in the adult NSCs.

Here, we provide evidence that paracrine signals released by neurons expressing E2F4DN increase the self-renewal, proliferation, and differentiation capacities of adult NSCs both in vivo and *in vitro*. These signals are mimicked by TrkB activation, while E2F4DN is able to upregulate the expression of NT4/5 in cortical neurons both *in vivo* and *in vitro*. We also show that blocking TrkB signaling prevents the improvement of NSCs proliferative and differentiation behavior by E2F4DN. These results confirm the capacity of neurons to release paracrine factors to act on ASN. In addition, they also stress the capacity of E2F4DN to act as a multifactorial therapeutic agent against AD: in this case due to the improvement of ASN.

## Methods

### Mice

Experimental procedures were conducted in 3–4-month-old transgenic and wild-type mice of both sexes on a C57BL/6J genetic background. Six months-old animals were also evaluated to analyze the pathological progression. The 5xFAD transgenic model, also maintained in C57BL/6J background, carries both human Aβ precursor protein 695 (APP_695_) containing the Swedish (K670N, M671L), Florida (I716V), and London (V717I) FAD mutations and human presenilin 1 (PS1) harbouring the M146L and L286V familial AD mutations under the control of the *Thy1* promoter (Tg6799 or 5xFAD mice). This murine AD model was purchased from The Jackson Laboratory (Bar Harbor, ME, USA) (strain #008,730). 5xFAD mice were crossed to obtain homozygote 5xFAD (h5xFAD) mice. Homozygosity was confirmed by quantitative PCR genotyping using specific primers of the human *APP* transgene: Fw 5’-AGGATGGTGATGAGGTAGAG-3’ and Rv 5’-CTGCTGTTGTAGGAACTCGA-3’. In addition, CD1 mice (Charles River; Wilmington, MA, USA) were used to obtain embryonic E17 cortical neurons. Mice were maintained in a 12-h light/dark cycle with free access to food and water *ad libitum*.

### NSCs culture and expansion

NSCs were isolated from 3-months-old mice. After cervical dislocation, brains were dissected out and the region containing the SVZ was isolated and cut into small fragments. The pieces were incubated with 0,025% Trypsin-EDTA (ThermoFisher Scientific) for 20 min at 37°C. After being centrifugated, tissue pellets were transferred to Dulbecco’s modified Eagle’s medium (DMEM)/F12 medium (1:1 v/v; ThermoFisher Scientific) and carefully triturated with a fire-polished Pasteur pipette to a single cell suspension. Isolated cells were collected by centrifugation (300 g for 10 min), suspended in DMEM/F12 medium containing 2 mM Glutamax (ThermoFisher Scientific), 1X B27 supplement without vitamin A (ThermoFisher Scientific), 1x Antibiotic/Antimycotic (ThermoFisher Scientific), 2 μg/ml heparin (Sigma-Aldrich) and supplemented with 20 ng/ml epidermal growth factor (EGF; ThermoFisher Scientific) and 10 ng/ml fibroblast growth factor (FGF; ThermoFisher Scientific) (NSCs medium) and maintained in a 95% air-5% CO_2_ humidified atmosphere at 37 °C (34). Neurospheres were allowed to develop for 10 days in these conditions. For culture expansion, primary neurospheres were disaggregated with 0.5 mM Accutase (Sigma-Aldrich) for 10 min and washed with NSCs medium without mitogens to generate single cells suspension. Then, 62.5 cell/µl were plated in fresh NSCs medium in a 95% air-5% CO_2_ humidified atmosphere at 37 °C and maintained for 7-8 passages. The number of primary and secondary neurospheres were counted manually by microscope. Neurosphere images were taken using a PAULA Smart Cell Imager (Leica), and their diameters were measured using the ImageJ software. Each NSCs culture was generated using both SVZs from one adult mouse.

### Cortical cultures and conditioned media from Ad5-transduced neurons

Cortical primary cultures were generated to obtain conditioned media. For that, E17 mouse embryos of CD1 genetic background were isolated and, after dissection, the brain was placed in Hank’s Balanced Salt Solution (HBSS; ThermoFisher Scientific) supplemented with 10 mM HEPES (ThermoFisher Scientific). The cortex was dissected, removing hippocampi and meninges, and fragmented in 5-6 pieces per animal. Cells were dissociated with 2.5 mg/ml (0.25%) Trypsin (ThermoFisher Scientific) and 1 mg/ml DNAse (Roche; Basel, Switzerland) for 18 min at 37°C. The samples were then washed in DMEM (ThermoFisher Scientific) supplemented with 10% inactivated horse serum (iHS; ThermoFisher Scientific) and 100 U/mL (1x) penicillin/streptomycin (P/S; ThermoFisher Scientific). The fragments were then carefully triturated with a fire-polished Pasteur pipette to a single cell suspension. Isolated cells were collected by centrifugation at 300 g for 10 min, suspended in DMEM/10% iHS plus 1x P/S and plated at 25,000 cells/cm^2^. After 1-2 h of plating, the medium was replaced with Neurobasal (ThermoFisher Scientific) supplemented with 2 mM Glutamax (ThermoFisher Scientific), 1x B27 Supplement (ThermoFisher Scientific) and 50 U/mL (0.5x) P/S. Half of the medium was replaced with fresh medium every other day. After 14 DIV, cortical cultures were transduced with either Ad5.E2F4DN or Ad5.GFP at 50 IU/cell. After 48h, culture media of the transduced cells (i.e. conditioned media) were collected. Conditional media containing 2 μg/ml heparin, 10 ng/ml FGF and 20 ng/ml EGF were used to grow NSCs. In case of NSCs differentiation assays with conditioned medium, only heparin and FGF was added.

### Self-renewal, proliferation and differentiation assays

To estimate self-renewal potential, NSCs were plated at low density (5 cell/µl) after Accutase disaggregation in the fresh NSCs medium in a 95% air-5% CO_2_ humidified atmosphere at 37°C. Four replicates for each culture were used, and the average value of the number of neurospheres was estimated. The proliferation activity of NSCs were determined by plating 6.25 cell/µl and counting the number of cells after 5 days growing during 7 consecutive passages at the time of the next passage. The cell growth curves were estimated by the ratio of cell production at each subculturing step multiplied by the number of cells at the previous point of the curve. The proliferation of the NSCs was also studied by plating 6.25 cell/µl at the same conditions as before. After 3 days, neurospheres were plated onto glass coverslips (VWR; Radnor, PA, USA) coated with 1X Matrigel (Corning, Glendale, AZ, USA) for 15 min. After allowing NSCs attachment, they were fixed for 15 min at 37°C with 2% PFA prepared in PBS. For the differentiation assay, 80,000 cells/cm^2^ were seeded in Matrigel-coated coverslips and incubated for 2 DIV in the NSC culture medium without EGF. Then, medium was replaced with fresh-medium without FGF and supplemented with 2% of FBS for 5 more days *in vitro* (7 DIV) to allow terminal differentiation (30). Cultures were fixed at 7 days of differentiation with 2% PFA in PBS for 15 min at 37°C. When appropriate, BDNF was blocked with 1µg/ml recombinant human TrkB-Fc chimera protein (R&D Systems; MN, Minnesota, USA). To block the BDNF pathway, 10 µM of the specific antagonists ANA-12 (MedChemExpress; Monmouth Junction, NJ, USA) or THX-B (MedChemExpress), which inhibit TrkB and p75^NTR^ signaling, respectively (31; 32), were used. As control, cultures were exposed to 1:1000 of DMSO (Sigma-Aldrich). For BDNF treatment, h5xFAD cultures were treated with 10 ng/ml recombinant BDNF (ThermoFisher Scientific) in self-renewal and proliferation assays, or 50 ng/ml BDNF in the differentiation assay, as previously described by Lozano-Ureña and Frade (2023).

### Adeno-associated viral (AAV) vectors

The mouse *E2f4* coding sequence (NCBI accession number NM_148952) with Thr249Ala/Thr251Ala mutations (*E2f4dn*) containing the Kozak sequence (ACCATGG), flanked by the human synapsin 1 (*hSyn*) promoter (33) and the mutant woodchuck hepatitits virus post-transcripcional regulatory element (WPRE) sequence described by Schambach et al. (2006) (34), which lacks tumorigenic capacity (WPRE3SL), was synthesized (GenScript) and cloned into the pcDNA 3.1 (+) plasmid (Invitrogen). The WPRE3SL cassette has the SV40 late polyadeylation signal sequence plus the upstream polyadenylation enhancer element repeated in tandem described by Schambach et al. (2007) (35). The expected sequence of the hSyn1-*E2f4dn*-WPRE3SL cassette was confirmed by Sanger sequencing. This cassette or the GFP sequence, flanked by ITR2 sequences, were subcloned into a transfer plasmid to generate recombinant single stranded AAV vectors referred to as AAV.PHP.eB.E2F4DN or AAV.PHP.eB.GFP [Viral Vector Production Unit (UPV), Barcelona, Spain]. These AAV vectors, which cross the blood brain barrier (40), were intravenously administered through the tail vein at 1.25 x 10^10^ vg/g body weight. One month later, animals were sacrificed for subsequent analyses.

### Adenoviral vectors

The coding sequence of human E2F4 (hE2F4) (NCBI accession number NM_001950) containing the Thr248Ala/Thr250Ala mutations, flanked by the *hSyn1* promoter and the WPRE3SL sequence, was synthesized (GenScript; Piscataway, NJ, USA) and cloned in the pcDNA3 plasmid (ThermoFisher Scientific; Waltham, MA, USA). Then, serotype 5 adenoviral vectors (Ad5) were generated as previously described by Miravet et al. (2014) (36). To this aim, recombinant plasmids were generated for either GFP or the hSyn1-*hE2f4dn*-WPRE3SL cassette flanked by inverted terminal repeats (ITRs) as well as all the genes necessary to generate the vector, except for E1, which codes for the E1A and E1B proteins. These latter proteins are expressed by the Q packaging cell line BI-HEK_293A, which was transfected with the recombinant plasmids. In this way, the neuronal *hSyn1* promoter was used to induce the expression of E2F4DN (Ad5.E2F4DN) or GFP (Ad5.GFP). To transduce the cells, 50 infectious units (IU)/cell were used to induce E2F4DN or GFP expression.

### EdU treatment

For *in vivo* treatment, one month after AAV vector injection, the thymidine analogue 5-ethynyl-2’-deoxyuridine (EdU) (Santa Cruz Biotechnology; Dallas, TX, USA) was administrated for 7 days at 0.2 mg/ml in drinking water containing 1% sucrose (Millipore; Burlington, MA, USA), which was replaced by fresh EdU/sucrose solution every other day. The consumed water was measured to determine differences in consumption between experimental groups. Twenty-one days after EdU withdrawal, animals were perfused and, after detecting EdU incorporation as described below, the number of EdU positive cells was estimated.

To identify NSCs in S-phase *in vitro*, cultures were incubated for 1h with 10 μM EdU in proliferating medium. Then, cells were fixed with 2% paraformaldehyde (PFA) (Sigma-Aldrich; Saint Louis, MO, USA) during 15 min at room temperature (RT), and EdU incorporation was evaluated by Click-iT reaction (see below).

### Immunohistochemistry and immunocytochemistry

For immunohistochemical staining, animals were transcardially perfused with 4% PFA in 0.1M phosphate buffer saline pH 7.4 (PBS). Vibratome (Leica; Wetzlar, Germany) was used to generate coronal brain sections (40 μm). Then, sections were washed in PBS and blocked for 1h with blocking solution (BS) prepared with PBS containing 0.2% Triton X-100 (Sigma-Aldrich), 10% Foetal bovine serum (FBS) (ThermoFisher Scientific), and 1% glycine (ThermoFisher Scientific). For E2F4 immunodetection, antigen retrieval was performed by incubating brain sections in a solution containing 10 mM Tris (Roche), 1 mM EDTA (Merck), and 0.05% Tween 20 (Sigma-Aldrich) at 80 °C for 30 min. Then, sections were incubated overnight at 4°C with primary antibodies (Table S1) in BS and, after washing with PBS, sections were incubated with secondary antibodies (Table S2) for 1h at RT in BS. 4′,6-Diamidine-2′-phenylindole dihydrochloride (DAPI) (Sigma-Aldrich) was used at 1μg/ml to counterstain cell nuclei. Sections were then washed three times with PBS and mounted with ImmunoSelect antifading mounting medium (Dianova; Hamburg, Germany). Images were captured at 40x magnification and analyzed with a SP5 confocal microscope (Leica) or Stellaris8 STED confocal (Leica). Cell populations were manually counted using the LAS AF Lite 2.6.0 program (Leica) and data was obtained as a percentage of positive cells relative to a subpopulation or to total DAPI labeled cells in the lateral ventricle wall using 5 sections from each animal, with 10 μm intervals between sections.

For immunocytochemical staining, fixed cells (2% PFA) were permeabilized and blocked for 1h with BS, and incubated at 4°C overnight with primary antibodies (Table S1) in the same solution. Cells were washed three times with PBS, and incubated for 1h at RT with secondary antibodies (Table S2) in BS. DAPI (1μg/ml) was used to counterstain cell nuclei. Cells were then washed three times with PBS and mounted with ImmunoSelect antifading mounting medium. Images were acquired at 20x or 40x magnification with a SP5 confocal microscope (Leica). Cell populations were manually counted using LAS AF Lite 2.6.0 program and data was obtained as a percentage of positive cells relative to total of DAPI labeled cells. At least 250 cells (in 10 randomized fields) for each independent culture were counted to performed the analysis.

### EdU detection

To detect EdU in previously immunostained vibratome sections, Click-iT reaction was performed. Previously, EGFP-induced tissue fluorescence was photobleached by light exposure prior to imaging (Fig. S2D). To eliminate the fluorescence of EGFP, first, the sections were permeabilized for 1h at RT with 0.1M phosphate buffer (PB) containing 0.5% Triton X-100 (0.5% Tx). Then, sections were incubated for 1h at RT and protected from light with PB containing 2 mM CuSO_4_ (Sigma-Aldrich), 6 μM Alexa Fluor® 488 azide (Life Technologies; Carlsbad, CA, USA) and 20 mg/ml sodium L-ascorbate (Sigma-Aldrich). Then, sections were washed three times with PBS. The number of EdU positive cells was analysed from LAS AF Lite 2.6.0 images and data was obtained as a percentage of positive cells relative to a subpopulation in the interest region. In the OB, EdU positive cells was estimated by quantifying the total number of positive cells normalized to the area. Five sections per animal, separated by 10 μm, were taken into account.

For EdU detection in previously immunostained cells, the samples were permeabilized for 20 min at RT with 0.5% Tx in PBS and, then, the Click-iT reaction was performed incubating for 30 min at RT with the reaction components protected from light. Then, cells were washed three times with PBS. The percentage of EdU positive cells was obtained relative to DAPI labeled cells.

### Gene expression analysis

To quantify mRNA levels, RNAs were extracted using RNAeasy mini kit (Qiagen; Hilden, Germany) including DNase treatment (Qiagen), following the manufacturer’s guidelines. For quantitative PCR (qPCR), 1 μg of total RNA was reverse transcribed using random primers and SuperScript IV Reverse Transcriptase (ThermoFisher Scientific), following the instructions of the manufacturer. Thermocycling was performed in a final volume of 15 μl, containing 1 μl of cDNA sample (diluted 1:10) and the reverse transcribed RNA was amplified by PCR with appropriate primers (see Table S3) with PrimePCR SYBR Green Assay (Cultek; San Fernando de Henares, Spain). qPCR was used to measure gene expression levels normalized to *Rpl27*, the expression of which did not differ between the groups. qPCR reactions were performed in a 7500real-time PCR equipment (Applied Biosystems; Waltham, MA, USA).

### Western Blotting analysis

Brain extracts from GFP or E2F4DN injected h5xFAD animals were obtained in cold extraction buffer (20 mM Tris-HCl Ph 6.8, 10 mM β-mercaptoethanol (Sigma-Aldrich), 1 mM EDTA (Merk), 1% Triton X-100, 1% SDS (Sigma-Aldrich) including 1x cOmplete Mini, EDTA-free, protease inhibitor cocktail (Roche)). The extracts were centrifuged for 10 min at 14,000xg (at 4 °C), and supernatants were boiled for 5 min in Laemli buffer. A total of 100 μg of protein, previously quantified using the Bradford assay (Protein Assay Dye Reagent Concentrate, BioRad), was loaded onto the gel. To fractionate the extracts, 4-15% NZY Tris-Glycine Precast gels (NYZtech) were used and transferred to Immobilon-FL membranes (Millipore). The membranes were first incubated for 1 h with Odyssey Blocking Buffer TBS (LI-COR), and the, incubated overnight at 4 °C with primary antibodies (Table S1). After washing membranes with TBS containing 0.1% Tween 20, they were incubated for 1 h at RT with 1:15,000 dilution of secondary antibodies (Table S2). The protein bands were visualized using the Odyssey Clx Infrared Imaging System (LI-COR).

### Olfactory habituation/dishabituation test

The olfactory habituation/dishabituation test was performed to evaluate odor detection and discrimination ability using a modified version of previously described protocols (37, 38). During the three days prior to testing, mice were handled daily to habituate them to the experimenter, the testing environment, and the testing cage, thereby minimizing stress-induced behavioral variability. On the day testing, mice were individually placed in the test cage and allowed to habituate to the environment for 15 min before the beginning of the experiment. Odor stimuli consisted of non-emotional value odor: distilled water (control) as vehicle, and 1% vanilla extract (Vanillin, Sigma-Aldrich) in distilled water. Fresh cotton swabs were prepared immediately before each trial by applying 50 μL of each odor solution and were presented through the wire cage lid. Vanilla odor was presented three consecutive times for 2 min per trial, with a 2 min inter-trial interval after the habituation with the vehicle odor. A new cotton swab was used for every trial to avoid odor contamination. During each presentation, the cumulative sniffing time was manually recorded. Sniffing behavior was defined as the animal orienting its nose toward the cotton swab within approximately 2 cm and actively investigating the odor source. Habituation was defined as a progressive decrease in sniffing time across repeated presentations of the same odor, whereas dishabituation was defined as an increase in sniffing time following the presentation of a novel odor.

### Statistical analysis

Statistical tests were performed using the GraphPad Prism Software, version 8.00 for Windows. Data were tested for normality with a Shapiro-Wilk test. For normal data, a *t*-test or one-way ANOVA followed by Tukey’s post-hoc test were used to compare two or more groups, respectively. When two independent variables were analyzed, a two-way ANOVA followed by Tukey’s post hoc test was performed. For data groups that did not pass normality, Mann-Whitney (for two categories) or Kruskal-Wallis (more than two categories) non-parametric tests were performed followed by post-hoc test. The presence of outlier values was evaluated by Grubb’s test. Values of *p*<0.05 were considered statistically significant. Data are presented as the mean ± standard error of the mean (SEM) and the number of independent cultures or animals (*n*) and *p*-values are indicated in the figures. Box and whisker plots show the mean (+), median (line in box), and maximum and minimum values (whiskers).

## Results

### E2F4DN-based gene therapy restores neurogenesis in h5xFAD mice

A gene therapy based on neuronal expression of E2F4DN has proved to act as a multifactorial therapeutic approach for AD (10), able to restore cognitive impairment in h5xFAD mice (39). To go deeper into the mechanisms involved in E2F4DN-dependent cognitive improvement, adult neurogenesis was studied at 3 and 6 months in the SVZ of WT and h5xFAD mice. This analysis demonstrated that neurogenesis is altered in the adult SVZ of h5xFAD animals. This was evidenced by the expression of the mRNAs encoding *Nestin*, *Ascl1* and *Dcx* (Fig. S1A, B), known markers associated with the progression of the adult neurogenesis process (40). These markers showed a statistically significant decrease of expression in 6-month-old h5xFAD mice, the stage when survival declines in this murine model of AD with aggravated phenotype (15), compared to WT mice (Fig. S1B. This suggests that ASN of h5xFAD mice is impaired at this later stage. However, downregulation of expression was already statistically significant for the *Nestin* gene at 3 months (Fig. S1A), in agreement with an evident reduction of Nestin protein in the SVZ of these animals (Fig. S1C). To further characterize how ASN is altered in h5xFAD mice, we evaluated by immunofluorescence the activity of the SVZ in h5xFAD vs WT mice at 6 months of age. The number of proliferating cells in the niche was estimated by immunostaining of Ki67, a marker of all cell cycle phases (41). This analysis showed a statistically significant decrease in the percentage of Ki67+ cells in h5xFAD mice compared to WT mice at this age (Fig. S1E).

To confirm that ASN is impaired in h5xFAD mice and verify that this process can be restored by neuronal expression of E2F4DN, ASN was studied in WT and h5xFAD mice treated at 3 months with an AAV.PHP.eB vector expressing E2F4DN (or GFP as a control) under the neuron-specific *hSyn1* promoter, and analyzed 1 month after injection (Fig. 1A). These vectors, which can cross the blood-brain barrier in C57BL/6J mice (42), were referred to as AAV.E2F4DN and AAV.GFP, respectively. An immunofluorescence study performed under control conditions (i.e. AAV.GFP-treated) revealed a decrease in the percentage of Ki67+ cells in the SVZ of h5xFAD mice compared to that of WT mice (Fig. 1B). Nevertheless, unlike what was observed in 6-month-old mice (see above), this decrease was not statistically significant, thus indicating that the ASN deficits are progressively associated with disease progression. Interestingly, the pool of GFAP+/SOX2+ NSCs and DCX+ neuroblasts was already decreased in the SVZ of 4-month-old control h5xFAD animals (Fig. 1C, D). Moreover, the percentage of active NSCs, identified as Ki67+/GFAP+/SOX2+ cells, was also reduced in control h5xFAD mice at this age (Fig. 1C), whereas no significant differences in the number of Ki67+ neuroblasts (Ki67+/DCX+ cells) was observed between h5xFAD and WT mice treated with the AAV.GFP vector (Fig. 1D). Finally, consistent to the reduction in the number of neuroblasts in the SVZ, a decrease in the percentage of the migrating DCX+ population through the RMS was detected in these control h5xFAD animals of 4 months of age (Fig. 2A).

**Fig. 1.**
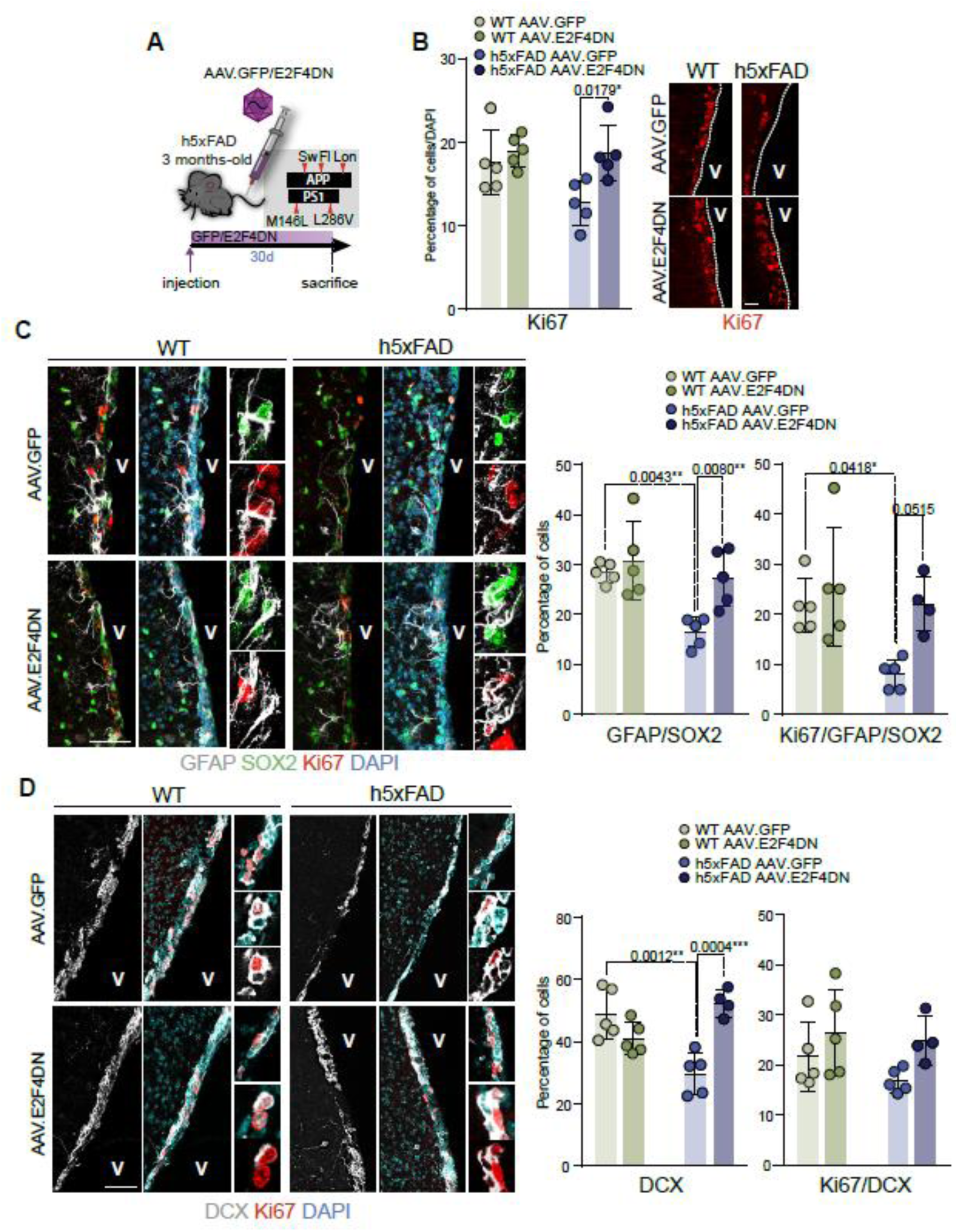
Neuronal expression of E2F4DN restores neurogenesis in h5xFAD SVZ. **A** Scheme showing the intravenous administration of AAV.E2F4DN and AAV.GFP through the tail vein in homozygous 5xFAD mice (h5xFAD), a transgenic model of Alzheimer’s disease carrying human APP and PS1 with 5 mutations triggering Familial AD (red head arrows). Tail vein injection was performed in 3-month-old animals using the AAV.PHP vector. Animals were sacrificed 1 month after the injection. **B** Percentage of Ki67 positive cells in the SVZ of WT and h5xFAD animals after 1 month of GFP (AAV.GFP) or E2F4DN (AAV.E2F4DN) administration (left panel). Right panel: immunohistochemistry for Ki67 (red) (two-way ANOVA followed by Tukey’s post-hoc test). **C** Immunohistochemistry (left panel) and percentage (right panel) of GFAP+/SOX2+ cells and Ki67+ cells within the GFAP+/SOX2+ (Ki67/GFAP/SOX2) population in the SVZ of 3-month-old WT and h5xFAD animals after 1 month of GFP (AAV.GFP) or E2F4DN (AAV.E2F4DN) injection (two-way ANOVA followed by Tukey’s post-hoc test). **D** Immunohistochemistry (left panel) and percentage (left panel) of DCX+ population (DCX) and Ki67+ cells within the DCX+ population (Ki67/DCX) in the SVZ of 3-month-old WT and h5xFAD animals after 1 month of GFP (AAV.GFP) or E2F4DN (AAV.E2F4DN) injection (two-way ANOVA followed by Tukey’s post-hoc test). DAPI was used to counterstain DNA. v: ventricle. Scale bars: 20 μm (B); 50 μm (C,D).

**Fig. 2.**
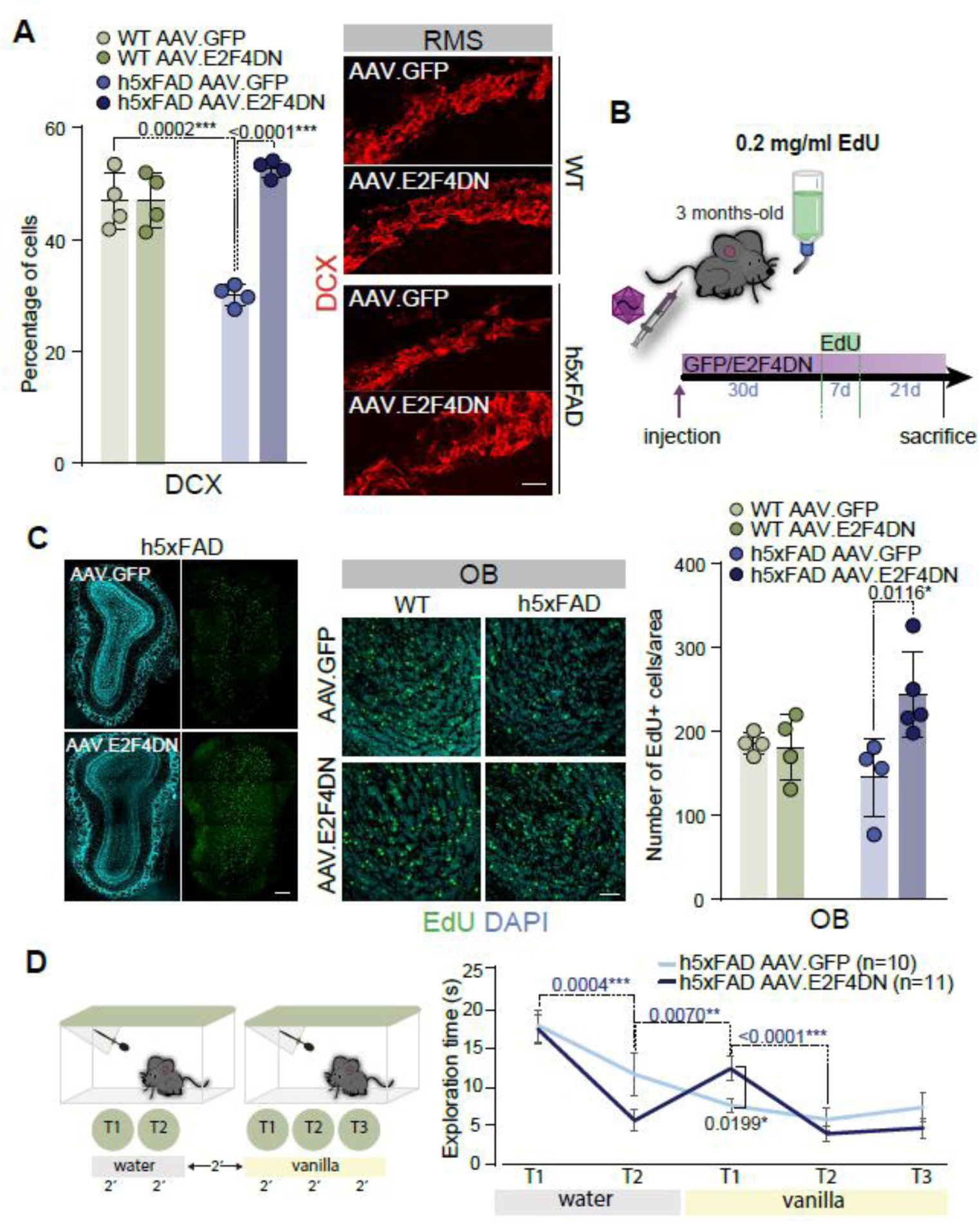
Neuronal expression of E2F4DN increases OB neuron replenishment in h5xFAD mice. **A** Percentage of DCX+ neuroblasts in the rostral migratory stream (RMS) of WT and h5xFAD animals after 1 month of GFP (AAV.GFP) or E2F4DN (AAV.E2F4DN) tail vein injection (left panel). Immunohistochemistry of DCX (red) is also shown (right panel) two-way ANOVA followed by Tukey’s post-hoc test). **B** Scheme illustrating the protocol used for EdU treatment in the drinking water in AAV.GFP-or AAV.E2F4DN-injected animals. Animals were sacrificed 21 days after one week of EdU treatment (one-way ANOVA followed by Tukey’s post-hoc test). **C** Confocal images of EdU (green) in the olfactory bulb (OB) of h5xFAD animals after AAV.GFP or AAV.E2F4DN tail vein injection and EdU treatment (left and middle panel). The number of EdU positive cells per μm^2^ in the OB of these animals is also indicated (right panel) (two-way ANOVA followed by Tukey’s post-hoc test). **D** Representation of olfactory habituation/dishabituation test (left panel). Time in seconds (s) of exploration time measured as time sniffing the odor in h5xFAD mice after one month of the GFP or E2F4DN treatment (dark blue p-values: AAV.E2F4DN comparisons, black p-value: E2F4DN vs GFP at vanilla T1; two-way ANOVA for repeated measures followed by Tukey’s post-hoc test). Scale bars: 20 μm (A); 200 μm (C; zoom: 50 μm).

To determine the potential role of E2F4DN in the regulation of ASN in h5xFAD animals, the percentage of Ki67+ proliferative cells was determined by immunofluorescence in the SVZ of 4-month-old h5xFAD and WT mice injected at 3 months with the AAV.E2F4DN vector (E2F4DN), in comparison to AAV.GFP-treated h5xFAD mice (Fig. 1B). As expected, qPCR demonstrated that the *E2f4dn* transcript was present in mice transduced with the AAV.E2F4DN vector (Fig. S2A, left). In accordance, expression of the E2F4 protein was detected in NeuN positive neurons of AAV.E2F4DN-treated mice (Fig. S2A, right). Although the reduction in the percentage of Ki67+ cells in h5xFAD mice showed a non-significant tendency, the injection of these animals with AAV.E2F4DN resulted in a statistically significant increase of the proliferative activity in the neurogenic niche (Fig. 1B). Specifically, the number of active (Ki67+/GFAP+/SOX2+) NSCs in the h5xFAD mice was significantly increased in E2F4DN treatment compared with control h5xFAD mice, reaching the WT values (Fig. 1C). In contrast, the percentage of Ki67+/DCX+ cells in h5xFAD mice treated with AAV.GFP or in E2F4DN-treated h5xFAD mice did not significantly differ from that observed in GFP-or E2F4DN-treated WT mice (Fig. 1D). Importantly, the percentages of GFAP+/SOX2+ NSCs and DCX+ neuroblasts were fully recovered in the SVZ of h5xFAD mice after injection with E2F4DN (Fig. 1C, D), indicating that neuronal expression of E2F4DN is a novel and efficient strategy to restore the neurogenic cell populations in the AD model.

### E2F4DN-based gene therapy increases olfactory bulb neuronal numbers and improves odor habituation/dishabituation in 5xFAD mice

Consistently with the pro-neurogenic effect of E2F4DN in the SVZ, a statistically significant increase of migrating DCX+ neuroblasts was detected at 4 months in the RMS of h5xFAD mice after the administration of AAV.E2F4DN one month earlier (Fig. 2A). To determine if the increase in NSCs and neuroblasts entails a consequent increment in the number of cells that reach and differentiate in the OB, 4-month-old h5xFAD and WT mice, injected at 3 months with AAV.GFP or AAV.E2F4DN, were treated with the thymidine analogue 5-ethynyl-2’-deoxyuridine (EdU) in the drinking water during 7 days, and sacrificed 21 days later (Fig. 2B). EdU is specifically incorporated during the S-phase of the cell cycle and diluted in each cell division, thus, only slowly proliferating NSCs, and cells that ceased to divide and underwent terminal differentiation soon after EdU incorporation can retain the nucleotide analogue 21 days after the EdU treatment (43). The consumed water was monitored as an internal control, resulting in similar values for all experimental conditions (Fig. S2B). This analysis showed that the number of EdU+ neurons reaching the OB (Fig. S2C) was significantly increased in the E2F4DN-injected h5xFAD mice, whereas no effect was observed in treated WT animals (Fig. 2C; Fig. S2D).

The decline in the number of NSCs and neuroblasts in h5xFAD mice does not seem to be due to a bias towards glial differentiation since the percentage of EdU+/OLIG2+ cells, identifying oligodendrocytic lineage, in the CC did not show alterations (Fig. S2E). Furthermore, the number of EdU+ astrocytes in the parenchyma was also unaltered in WT and h5xFAD animals (Fig. S2F), while the injection of E2F4DN did not affect the number of glial cells (Fig. S2E, F).

Alterations in ASN result in changes in odorant discrimination which can be assayed in habituation-dishabituation behavioral paradigms (37, 38). To evaluate the physiological relevance of improving ASN in h5xFAD animals, the olfactory habituation/dishabituation test was performed in h5xFAD animals treated with GFP or E2F4DN (Fig. 2D). The increment in the number of new neurons achieving the OB in the E2F4DN-treated h5xFAD animals was accompanied with an improvement in the olfactory behavior. Animals were first exposed to water and subsequently to the vanilla odor. E2F4DN-treated animals showed increased exploration of the vanilla odor, followed by a progressive decrease across the three consecutive trials, indicating habituation. In contrast, GFP-injected animals did not show significant changes in the exploration time of the vanilla odor across the three consecutive trials (Fig. 2D), indicating absence of discrimination between both odors. Consistently, the time spent sniffing the vanilla in the trial 1 was significantly lower in GFP treated h5xFAD animals compared to E2F4DN group (Fig. 2D).

### Self-renewal and proliferative capacities are impaired in NSCs isolated from the SVZ of adult h5xFAD mice

The alterations in the GFAP+/SOX2+ NSCs and the DCX+ neuroblasts that are observed in the adult SVZ of h5xFAD mice (Fig. 1C, D) suggest that a decreased ability of NSCs to proliferate and differentiate is among the perturbations occurring in AD. To test whether h5xFAD-derived NSCs maintain these properties *in vitro*, NSCs were isolated from the SVZ of 3-month-old WT and h5xFAD mice, and then cultured as neurospheres under proliferating conditions (43), and the number of primary neurospheres generated was estimated. This assay resulted in fewer number of neurospheres in h5xFAD compared to WT cultures (Fig. 3A). To confirm alterations in the self-renewal capacity of the h5xFAD-derived cultures, primary neurospheres were dissociated into single cells and plated at low density. h5xFAD NSCs also showed a reduction in the number of secondary neurospheres (Fig. 3A). Moreover, the diameter of the primary and secondary neurospheres was also reduced in h5xFAD compared to WT cultures (Fig. 3B), suggesting the affectation of the NSCs proliferation ability as well. To assess the proliferative capacity of the NSCs, neurospheres were dissociated into single cells and plated at stablished cell density for several passages, evaluating the number of cells after 5 days of each dissociation. The resulting growth curve showed an impaired expansion of the h5xFAD cultures compared to WT NSCs (Fig. 3C), supporting a defect in the proliferative capacity of the NSCs in the AD context. The number of proliferating NSCs was also evaluated by immunocytochemistry for Ki67. Thus, a reduction in the percentage of Ki67+ cells was also observed in h5xFAD cultures (Fig. 3D), confirming the impaired proliferation ability of the h5xFAD NSCs.

**Fig. 3.**
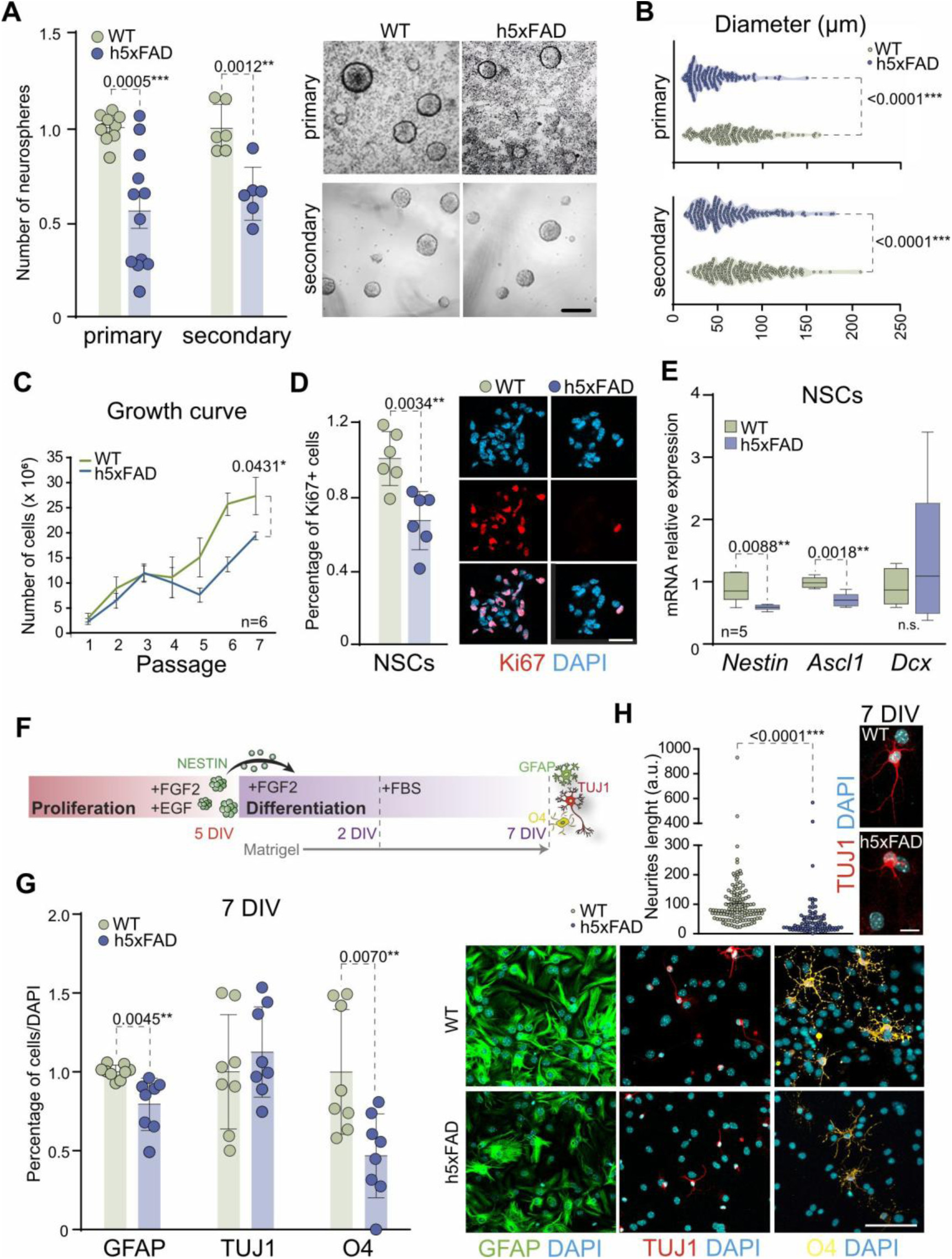
Self-renewal, proliferation and differentiation capacities are impaired in h5xFAD NSCs. **A** Number of primary and secondary neurospheres in WT and h5xFAD cultures of 3-month-old mice (left panel). Representative images are also shown (right panel) (two-tailed Student’s t-test). **B** Diameter of the primary and secondary neurospheres in the WT and h5xFAD cultures (two-tailed Student’s t-test). **C** Growth curve of WT and h5xFAD cultures. **D** Percentage of Ki67 NSCs in WT and h5xFAD cultures (left panel). Representative images are also shown (right panel) (two-tailed Student’s t-test). **E** Gene expression of *Nestin*, *Ascl1* and *Dcx* in the NSCs of the SVZ from WT and h5xFAD mice. *Rpl27* was used to normalize data (two-tailed Student’s t-test). **F** Scheme of NSCs differentiation protocol. Proliferative neurospheres maintained for 5 DIV are disaggregated and NSCs are then plated in Matrigel-coated dishes in the presence of FGF during 2 DIV. Afterwards, FGF is removed and 2% FBS is added to the medium during 5 additional days. **G** Percentage of GFAP astrocytes, TUJ1 neurons and O4 oligondendrocytes in WT and h5xFAD cultures after 2+5 days (7 DIV) under differentiation conditions (left panel) (two-tailed Student’s t-test and Mann-Whitney non-parametric test). Representative immunofluorescence images are shown (right panel). **H** Neurite length in TUJ1 positive cells after 7 DIV under differentiation conditions of WT and h5xFAD cultures (Mann-Whitney non-parametric test). DAPI was used to counterstain DNA. Scale bars: 200 μm (A); 20 μm (D); 50 μm (G); 10 μm (H).

Consistently with the alterations in the expression of genes associated with the process of ASN in the SVZ of h5xFAD mice (Fig. S1A), the mRNA levels of the NSCs/progenitor markers *Nestin* and *Ascl1* were also reduced in the NSCs isolated from these animals (Fig. 3E). Immunofluorescence detection for Nestin protein also showed lower levels in the h5xFAD cultures (Fig. S1D). As expected, no changes in mRNA levels of more differentiating markers such as the neuroblast marker *Dcx* was observed in the NSCs cultures (Fig. 3E).

### h5xFAD NSCs fail to undergo terminal differentiation *in vitro*

To investigate the possible alteration of the multipotencial capacity of h5xFAD NSCs *in vitro*, we measured the percentage of the GFAP+ cells (i.e. astrocytes), the βIII-tubulin (TUJ1+) cells (i.e. neurons), and the O4+ cells (i.e. oligodendrocytes) that were present after 7 DIV under NSCs differentiation promoting conditions (Fig. 3F, G). Whereas no significant differences were detected in the percentage of TUJ1+ neurons between WT and h5xFAD cultures, a reduction in the percentage of both astrocytes and oligodendrocytes was observed in the h5xFAD-derived cultures (Fig. 3G). Despite the lack of differences in the percentage of cells expressing the immature neuronal marker TUJ1, measurement of the neurite length indicated that the new-formed neurons observed in the h5xFAD-derived cultures showed shorter prolongations compared to the WT cultures (Fig. 3H), suggesting alterations in neuronal maturation in the AD model.

The analysis at an early stage of differentiation (2 DIV) showed that the impairment in NSCs commitment already started at this time point. The expression of *Olig2* at 2 DIV of differentiation was significantly decreased in h5xFAD compared to WT cultures (Fig. S3A), agreeing with the impairment in oligodendrocytes differentiation at 7 DIV (Fig. 3G). The expression of the astrocytic marker *S100β* and the neuronal marker *βIII-tubulin* was not affected at 2 DIV under differentiation conditions (Fig. S3A). However, the expression of *S100β* was statistically significant lower in the h5xFAD compared to WT cultures at 7 DIV of differentiation (Fig. S3B), which is consistent with the impairment in astrocytic differentiation observed in the h5xFAD cultures (Fig. 3G). No significant changes were observed in the expression of *βIII-tubulin* at 7 DIV of differentiation in h5xFAD cultures, although a clear tendency to reduced levels of this neuron-specific transcript was detected in the h5xFAD-derived cells (Fig. S3B).

### Conditioned medium by neuronal E2F4DN promotes self-renewal and proliferation of h5xFAD NSCs *in vitro*

To understand the mechanisms of neuronal E2F4DN in the promotion of neurogenesis in the SVZ of h5xFAD mice, WT and h5xFAD NSCs were cultured in medium conditioned by neurons transduced with Ad5.GFP (NSCs-WT^GFP^ and NSCs-h5xFAD^GFP^, respectively) or Ad5.E2F4DN (NSCs-WT^E2F4DN^ and NSCs-h5xFAD^E2F4DN^, respectively) for 48 h (Fig. 4A). As expected, the Ad5.E2F4DN-transduced neurons expressed the *E2f4dn*-specific transcript (Fig. S4A). Single cell suspensions of both WT- and h5xFAD-derived NSCs were grown during 5 days in E2F4DN-or GFP-conditioned media after adding EGF and FGF to induce proliferative conditions (Fig. 4A). The number of neurospheres generated after 5 DIV was significantly decreased in NSCs-h5xFAD^GFP^ as compared to NSCs-WT^GFP^ or NSCs-WT^E2F4DN^, while their number was restored in NSCs-h5xFAD^E2F4DN^ (Fig. 4B), indicating that E2F4DN-expressing neurons stimulate self-renewal capacity on the NSCs in a cell-to-cell independent mechanism. This finding was consistent with an increase in the expression of *Nestin* in NSCs-h5xFAD^E2F4DN^ compared to NSCs-h5xFAD^GFP^, whereas no recovery of *Ascl1* expression was observed (Fig. 4C).

**Fig. 4.**
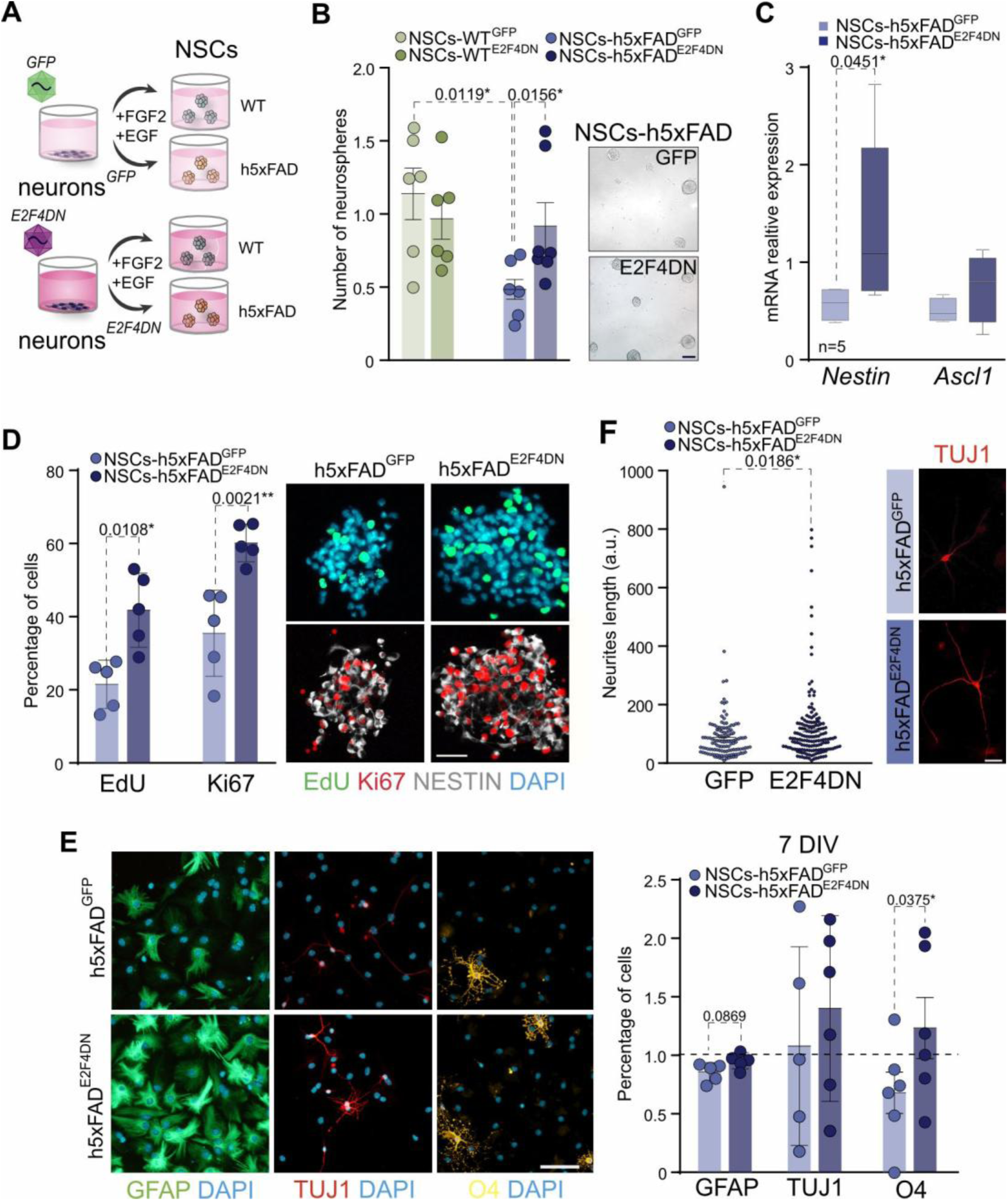
E2F4DN-conditioned medium restores h5xFAD NSCs behavior**. A** Scheme of the assay for NSCs conditioning with cortical cultures transduced with AAV.GFP (GFP) or AAV.E2F4DN (E2F4DN). **B** Number of neurospheres in WT- and h5xFAD-derived cultures after GFP or E2F4DN-conditioning medium (left panel). Representative images are shown (right panel) (Kruskal-Wallis non-parametric test). **C** Gene expression of *Nestin* and *Ascl1* in h5xFAD NSCs after 5 DIV in the presence of GFP-or E2F4DN-conditioned medium. *Rpl27* was used for normalization (two-tailed Student’s t-test). **D** Percentage of EdU and Ki67 cells in h5xFAD NSC cultures after 5 DIV in the presence of GFP-or E2F4DN-conditioned medium (left panel). Representative images are shown (right panel) (two-tailed Student’s t-test). **E** Percentage of GFAP, TUJ1 and O4 positive cells in h5xFAD cultures after 7 DIV in conditioned medium with GFP or E2F4DN-conditioned medium under differentiation conditions (right panel) (two-tailed Student’s t-test). Representative images are shown (left panel). DAPI was used to counterstain DNA. **F** Neurite length in TUJ1 positive cells present in h5xFAD NSCs cultured for 7 DIV under differentiation conditions with GFP-or E2F4DN-conditioned medium (left panel) (Mann-Whitney non-parametric test). Representative images are shown (right panel). Scale bars: 100 μm (B); 20 μm (D,E); 50 μm (F).

The NSCs proliferation rate was estimated as the percentage of cells in S-phase, based on a short pulse of EdU, and the percentage of Ki67+ cells. NSCs-h5xFAD^E2F4DN^ showed an increase of proliferation markers compared with NSCs-h5xFAD^GFP^ (Fig. 4D), indicating a proliferative effect of neuronal E2F4DN on the h5xFAD-derived NSCs in a cell-to-cell independent manner, according to the *in vivo* effect. Moreover, the diameter of the neurospheres was increased in the cultures generated from the SVZ of AAV.E2F4DN-injected h5xFAD animals compared to control h5xFAD animals injected with AAV.GFP (Fig. S4B). Altogether, these results demonstrate the potential role of neuronal expression of E2F4DN to recover the proliferation deficits of NSCs in this AD model.

### Conditioned medium by E2F4DN-expressing neurons promotes commitment of h5xFAD NSCs to oligodendrocytes and morphological differentiation of neurons *in vitro*

In addition to impaired self-renewal and proliferation, h5xFAD NSCs also showed alterations in their differentiation capacity (see above). To reveal the implications of neuronal expression of E2F4DN in NSCs differentiation, h5xFAD NSCs were cultured for 2 DIV in Matrigel-coated glasses in the presence of either E2F4DN-or GFP-conditioned media containing FGF. Then, these media were replaced by conditioned media supplemented with 2% FBS during 5 additional days. After these 7 days under differentiating conditions, the effect of the E2F4DN-or GFP-conditioned media on the percentage of GFAP+ astrocytes, TUJ1+ neurons and O4+ oligodendrocytes was determined. The slight decrease observed in the percentage of astrocytes in h5xFAD cultures compared to WT (Fig. 3G) was not statistically recovered after conditioning with E2F4DN (Fig. 4E). Instead, the percentage of oligodendrocytes was significantly recovered in cultures conditioned by neurons transduced with E2F4DN (Fig. 4E). No alterations in the percentage of neurons was detected in the h5xFAD cultures conditioned with GFP-or E2F4DN-conditioned medium (Fig. 4E). However, TUJ1+ cells showed a longer neurite length when h5xFAD NSCs were differentiated with E2F4DN-conditioned medium compared to GFP-conditioned medium (Fig. 4F). The positive effect of the conditional medium of E2F4DN-expressing neurons in the behavior of cultured NSCs suggests the release of a neuronal factor with the ability to promote self-renewal, proliferation and differentiation of adult NSCs.

### E2F4DN-conditioned medium recovers NSCs behavior through TrkB signaling

BDNF is a neurotrophin that promotes the self-renewal, proliferation, and differentiation of neural stem cells through activation of TrkB and TrkB/p75^NTR^ signaling pathways (26), exerting effects similar to those of E2F4DN-conditioned medium in these cells. Both receptors (TrkB and p75^NTR^) are also activated with similar affinity by NT4/5, another member of the neurotrophin family (44), whose encoding mRNA (*Ntf5*) is significantly increased in neurons expressing E2F4DN (Fig. 5A). *Bdnf* mRNA, which encodes BDNF, showed a non-significant tendency toward upregulation in cortical cultures transduced with Ad5.E2F4DN compared with Ad5.GFP-transduced cultures. (Fig. 5A). Collectively, these findings indicate that TrkB activation may be responsible for the effects of E2F4DN-conditioned medium on the proliferation, self-renewal, and differentiation of h5xFAD NSCs.

**Fig. 5.**
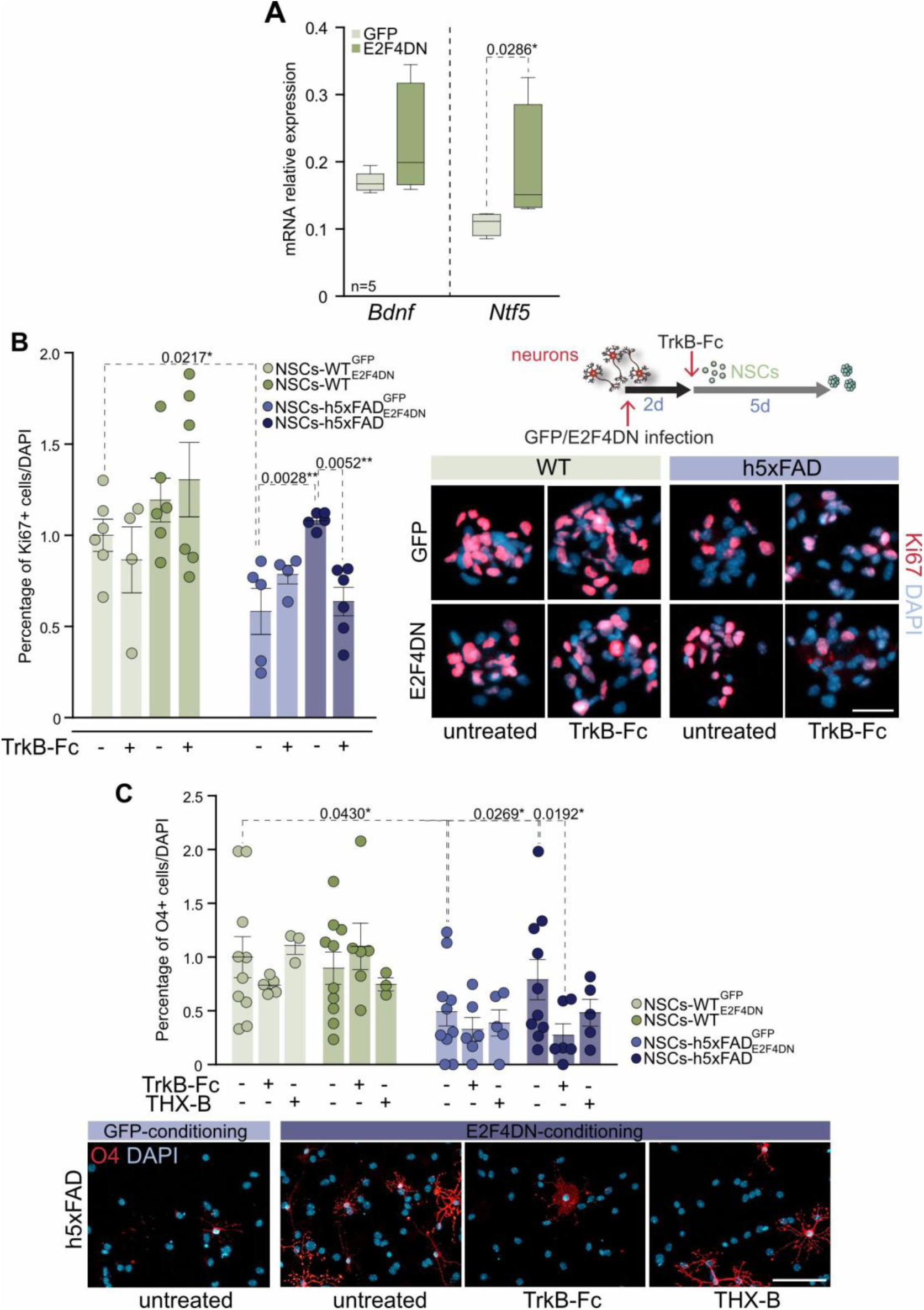
Blockage of TrkB signaling reverts the effect of E2F4DN-conditioned medium in h5xFAD NSCs. **A** Gene expression of *Bdnf* and *Ntf5* in cortical cultures from E17 mouse embryos used to generate the conditioned media. *Rpl27* expression was used for normalization (Mann-Whitney non-parametric test). **B** Percentage of Ki67 positive NSCs grown with GFP-or E2F4DN-conditioned medium in the presence or absence of the TrkB blocker TrkB-Fc (left panel). Right panel: scheme of the methodology (right upper panel) and representative immunofluorescence images (right bottom panel). **C** Percentage of O4 positive cells after NSCs differentiation in GFP-or E2F4DN-conditioned medium in the presence or absence of either TrkB-Fc or the p75^NTR^ antagonist THX-B (upper panel) (one-way ANOVA followed by Tukey’s post-hoc test and). Representative images are shown (bottom panel). DAPI was used to counterstain DNA. Scale bars: 20 μm (B); 50 μm (C).

To examine the involvement of TrkB signaling in the E2F4DN-induced recovery of h5xFAD NSC function, recombinant BDNF (10 ng/ml) was added to the cultures to activate TrkB signaling (26), and proliferative and self-renewal capacities were analyzed. The activation of TrkB increased the number of h5xFAD neurospheres compared to control h5xFAD NSCs (Fig. S5A, left panel). Moreover, the diameter of the neurospheres were significantly higher (Fig. S5A, right panel), and the percentage of Ki67+ proliferative NSCs was increased in the TrkB-activated h5xFAD cultures compared to vehicle treatment (Fig. S5B). To study the effect of TrkB together with its accessory receptor p75^NTR^ (26) in the differentiation capacity of h5xFAD NSCs, these cells were differentiated as described above in the presence or absence of 50 ng/ml BDNF to activate the TrkB/p75^NTR^ complex. This manipulation resulted in an increased proportion of neurons and oligodendrocytes at the expense of the astrocytic population (Fig. S5C). Taken together, these results indicate that the effects of E2F4DN-conditioned medium on the self-renewal, proliferation, and differentiation of h5xFAD NSCs are likely mediated by TrkB signaling.

To confirm the involvement of TrkB in E2F4DN-mediated promotion of NSC proliferation, a fusion protein comprising the Fc domain of human IgG and the extracellular domain of TrkB (TrkB-Fc receptor body; 26), which inhibits both NT4/5 and BDNF signaling (45), was added to media conditioned by GFP-or E2F4DN-expressing neurons. TrkB-Fc treatment prevented the increase in the percentage of Ki67+ cells observed in h5xFAD NSCs exposed to E2F4DN-conditioned medium, whereas no effect was detected in h5xFAD NSCs exposed to GFP-conditioned medium or in WT NSCs under either condition (Fig. 5B).

To further confirm that the increase of proliferation observed in NSCs-h5xFAD^E2F4DN^ relies on TrkB signaling, an antagonist of TrkB (ANA-12) (26) was added to the E2F4DN conditioned medium. While the blockage of the TrkB pathway with ANA-12 had no effect in NSCs-WT^GFP^ or NSCs-h5xFAD^GFP^ cultures, NSCs-h5xFAD^E2F4DN^ cultures showed a statistically significant decrease in the percentage of Ki67+ cells in the presence of this TrkB antagonist (Fig. S5D). These results confirm the implication of the TrkB pathway in the promotion of h5xFAD NSCs proliferation by the E2F4DN-conditioned medium.

The role of TrkB signaling in driving oligodendrocytic differentiation in h5xFAD cultures in response to E2F4DN-conditioned medium was also investigated. As previously, TrkB-Fc was added in the conditioned media to block the TrkB signaling and the percentage of oligodendrocytes was estimated in WT and h5xFAD NSC differentiative cultures. This manipulation led to a reduction in the percentage of O4+ oligodendrocytes in h5xFAD cultures grown in E2F4DN-conditioned medium containing TrkB-Fc compared to untreated E2F4DN-conditioned h5xFAD cultures (Fig. 5C). No effects were observed in WT cultures regardless of the presence of TrkB-Fc (Fig. 5C).

As indicated above, p75^NTR^, the common receptor for all neurotrophins, including BDNF and NT4/5, has been previously implicated in NSC differentiation toward the oligodendrocytic lineage as a co-receptor in TrkB signaling (26). Therefore, to study the implication of p75^NTR^ signaling in the promotion of O4+ oligodendrocyte differentiation by E2F4DN, the p75^NTR^ antagonist THX-B (26) was used to block the p75^NTR^ pathway. Although a reduction was detected in E2F4DN-conditioned h5xFAD cultures in the presence of the p75^NTR^ antagonist, this reduction did not reach the statistically significance (Fig. 5C).

### E2F4DN promotes the expression and activation of TrkB in h5xFAD animals

The results obtained from the use of the TrkB inhibitors in the NSCs cultures confirm the implication of TrkB signaling in the regulation of proliferation, self-renewal and differentiation of NSCs through E2F4DN treatment. This is consistent with the observed significant upregulation of *Ntf5* in Ad5.E2F4DN-transduced neurons (see above). Moreover, the administration of the AAV.E2F4DN vector *in vivo* also increased the expression of *Ntf5* in the brain of h5xFAD animals compared to GFP-treated h5xFAD animals, whereas no effect was detected in *Bdnf* expression (Fig. 6A).

**Fig. 6.**
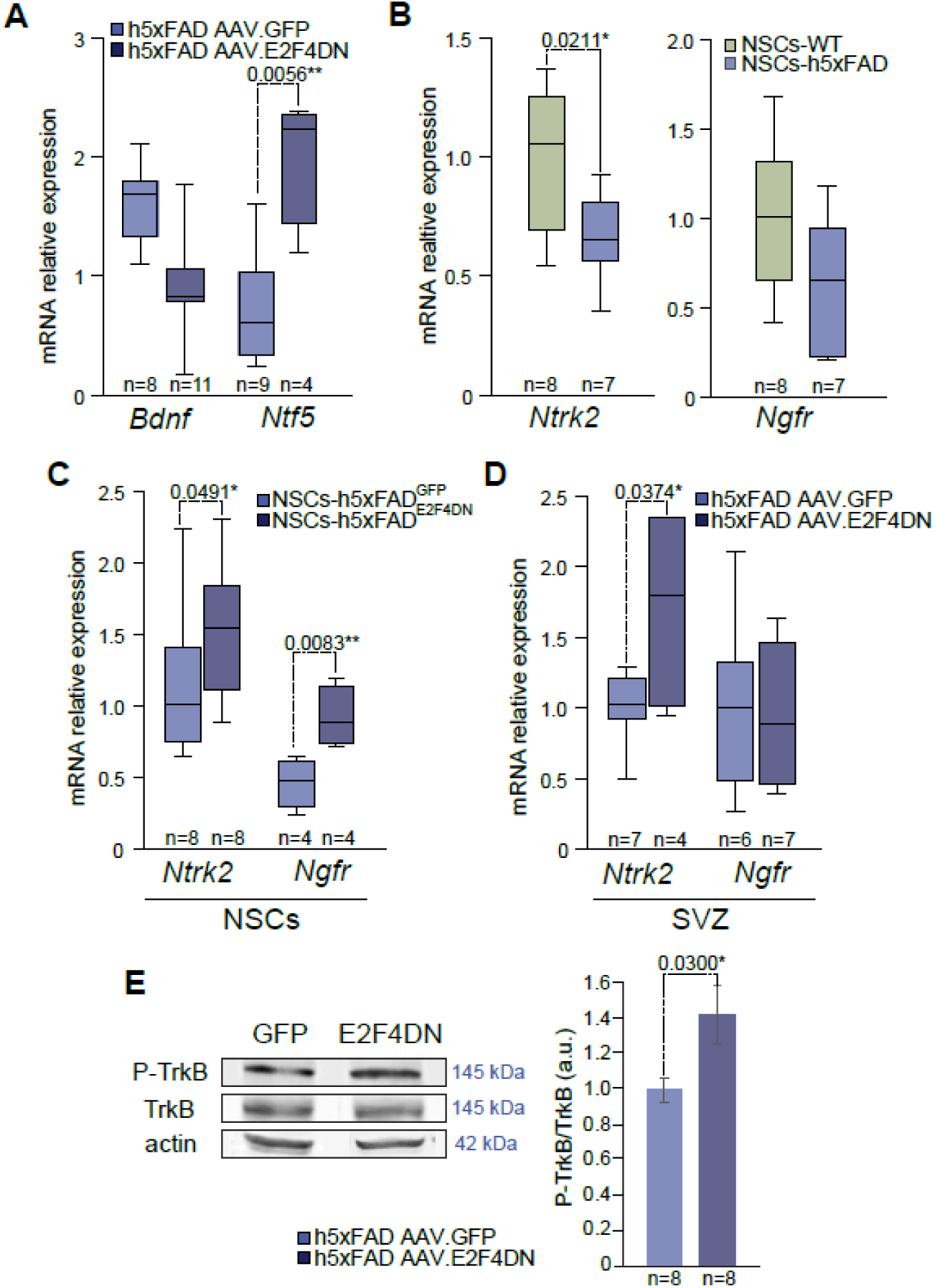
E2F4DN promotes the transcription of *Ntf5* and its receptors. **A** Gene expression of *Bdnf* and *Ntf5* in the cerebral cortex of h5xFAD animals treated with AAV.GFP or AAV.E2F4DN (two-tailed Student’s t-test). **B** Gene expression of *Ntrk2* and *Ngfr* in WT and h5xFAD NSCs (two-tailed Student’s t-test). **C** Gene expression of *Ntrk2* and *Ngfr* in h5xFAD NSCs grown with GFP-or E2F4DN-conditioned medium (two-tailed Student’s t-test). **D** Gene expression of *Ntrk2* and *Ngfr* in the SVZ of h5xFAD animals treated with AAV.GFP or AAV.E2F4DN (two-tailed Student’s t-test). **E** Quantification of the western blot for P-TrkB and TrkB in the cerebral cortex of GFP and E2F4DN-treated h5xFAD animals (two-tailed Student’s t-test). Actin was used as a loading marker. Representative images for the western blot are shown. *Rpl27* was used to normalize data.

The expression of TrkB-encoding mRNA (*Ntrk2*) was assessed in NSCs as a readout of TrkB signaling (26) to further confirm the involvement of this receptor in the signaling triggered by E2F4DN-conditioned medium. This analysis revealed that *Ntrk2* expression is reduced in NSCs and in the SVZ of h5xFAD mice compared to wild-type controls (Fig. 6B, left panel; Fig. S6A). These observations suggest that TrkB activity is reduced in h5xFAD mice, leading to a corresponding decline in the proliferation, self-renewal, and differentiation of these cells (26), as previously shown (Fig. 3). As expected, the expression of *Ntrk2* was significantly upregulated in the h5xFAD NSCs treated with E2F4DN-conditioned medium and in the h5xFAD SVZ of E2F4DN-treated animals (Fig. 6C, D), suggesting that TrkB signaling is activated by the E2F4DN-expressing neurons in the neurogenic niche, thus leading to an increase of expression of *Ntrk2* in the AD context.

To further demonstrate the capacity of E2F4DN to activate the TrkB signaling, the levels of active phospho-TrkB was determined by western blot in the cerebral cortex of E2F4DN-treated mice, compared to GFP-treated h5xFAD animals (Fig. 6E). E2F4DN expression induced an increase of TrkB phosphorylation at Tyr516 (Fig. 6E), a residue known to participate in TrkB signaling (46). Thus, treatment of h5xFAD animals with E2F4DN demonstrated *in vivo* activation of TrkB.

The analysis of the expression of the gene encoding p75^NTR^ (*Ngfr*), the common neurotrophin receptor which also becomes upregulated in NSCs once it becomes activated (26), showed a non-significant tendency to become reduced in h5xFAD NSCs compared to WT NSCs (Fig. 6B, right panel) while, in the h5xFAD SVZ, it was significantly reduced at both 3 and 6 months (Fig. S6B). This finding is consistent with a reduced activity of p75^NTR^, and consequently, a decline in the differentiation (26). Moreover, the expression of *Ngfr* was significantly increased in the h5xFAD NSCs conditioned with E2F4DN-derived medium compared to GFP medium (Fig. 6C). These results suggest that TrkB signaling is potentiated by p75^NTR^ in the E2F4DN-mediated recovery of adult NSC differentiation.

## Discussion

Despite the efforts to understand the implications of adult neurogenesis in AD, the debate still remains open (40). Whereas most studies support a decline in adult neurogenesis associated with AD, some others have found no changes or even an increase of neurogenic activity in this disease (47). However, olfactory impairment has been proposed as a neuropathological biomarker of AD (4), being the most prominent sensory system affected in this disease (48, 49). This alteration suggests that ASN could be altered early in the pathology as this process is known to replace interneurons in the OB during adulthood. In this study we have demonstrated that the capacity of the SVZ to generate new neurons is altered in h5xFAD mice compared to WT mice already at the early stage of 3-4 months. Although the expression of *Ascl1* and *Dcx*, or the percentage of Ki67 were no altered in the SVZ at 3 months, the analysis of these parameters at 6 months showed a dysregulation in the h5xFAD animals, making evident the deterioration of ASN throughout the disease progression. Our results are consistent with previous analyses performed in other mouse models of AD with lesser aggravated phenotype (1, 50–56) or in AD patients (57, 58).

E2F4 has been shown to fulfill multiple functions in a context-dependent manner (59). Despite the extensive study of the role of E2F4 in regulating the cell cycle (60, 61), the implication of this factor in tissue homeostasis is gaining increasing importance (10, 59, 62). Indeed, the modification of E2F4, preventing the phosphorylation of its Thr249/Thr251 conserved motif, has emerged as a promising therapy against AD thanks to its multifactorial action, as it reduces Aβ accumulation, attenuates neuroinflammation, and restores cognitive impairment (13, 39). In this study, we have shown that neuronal expression of E2F4DN is able to increase the number of NSCs and neuroblasts in the SVZ, increasing the number of neuroblasts migrating through the RMS and, therefore, the number of new neurons reaching the OB in h5xFAD mice, restoring the neurogenic populations. The recovery of proliferating NSCs (i.e. Ki67+/GFAP+/SOX2+ cells), which were present at lower levels in h5xFAD mice, suggests that neuronal expression of E2F4DN could be favoring NSCs activation, which give rise to more neuroblasts that migrate to the OB. In addition, a promotion of neuronal differentiation by E2F4DN was also observed in E2F4DN-condicioned medium, as this treatment allowed h5xFAD neuroblasts to generate neurites with longer length under differentiating-promoting conditions. Importantly, the enhanced ASN induced by E2F4DN in h5xFAD mice correlated with improved olfactory performance in the habituation/dishabituation test (36, 37), a finding consistent with previous studies on the role of ASN in odor discrimination (63–65). SVZ-derived NSCs respond dynamically to different physiological and pathological cues, generating not only neurons but also astrocytes and oligodendrocytes (66–68). Therefore, restoration of the SVZ NSC pool may have broader implications beyond enhancing neurogenesis, potentially contributing to endogenous repair mechanisms engaged in response to the multiple forms of neural damage occurring throughout disease progression.

In this study, the mechanism employed by E2F4DN to regulate ASN in h5xFAD mice was investigated using adult NSCs isolated from the SVZ and cultured in the presence of conditioned medium from GFP-or E2F4DN-transduced cortical neurons. As observed *in vivo*, the proliferation and differentiation capacity of h5xFAD NSCs was impaired compared to those from WT under control conditions. Therefore, NSCs maintain long-lasting modifications, likely of epigenetic nature, that can be kept after isolation. Indeed, epigenetic regulators have also emerged as key players in the regulation of NSCs, neural progenitor cells, and their differentiated progeny via modifications that include DNA methylation, histone modification, chromatin remodeling, and RNA-mediated transcriptional regulation (69, 70), with implications in AD (71).

Conditioned medium from E2F4DN-expressing neurons restored the self-renewal and proliferative capacity of h5xFAD NSCs as both the number and size of h5xFAD-derived neurospheres was increased under these conditions. These results are consistent with the role of extrinsic neuron-derived signals in adult neurogenesis, including neurotransmitters (72, 73), growth factors (74, 75), and hormones (76). The ability of conditioned medium from E2F4DN-expressing neurons to restore the self-renewal and proliferative capacity of h5xFAD NSCs is consistent with findings from (18). These authors showed that neuronal deletion of the p38^MAPK^ member family p38α, which is implicated in E2F4 phosphorylation, delays the age-associated exhaustion of NSCs. Therefore, neuronal deletion of p38α would reduce E2F4 phosphorylation, equivalent to neuronal E2F4DN expression, increasing the NSC population. In addition, E2F4DN-conditioned medium was able to increase the percentage of oligodendrocytes and the length of the neurites of NSC-differentiated neurons in vitro. Thus, under differentiation-promoting conditions, E2F4DN is able to favor NSCs commitment. Altogether, these results suggest that the expression of E2F4DN in neurons promotes the release of paracrine factor(s) capable of modulating h5xFAD NSCs behavior.

As E2F4DN-condioned medium, TrkB signaling promotes both NSCs proliferation and differentiation (26, 77), the latter together with the p75^NTR^ receptor (26). These two receptors that can be activated by NT4/5 (28, 78), and this latter neurotrophin can be endogenously expressed in the cerebral cortex (79). Our study has demonstrated that the expression of the gene encoding NT4/5 is significantly upregulated in cortical neurons expressing E2F4DN and in the cortex of h5xFAD mice treated with E2F4DN. The capacity of E2F4DN to modulate the expression of this neurotrophin is consistent with the presence of 10 JASPAR E2F4 motifs within the promoter of the mouse *Ntf5* gene (p<0.001) (Eukaryotic Promoter Database; https://epd.expasy.org/epd/). In this regard, our results suggest that NT4/5 participates in the regulation of h5xFAD neural stem cell behavior via E2F4DN signaling through TrkB, thereby favoring proliferation, self-renewal, and differentiation. Nevertheless, the presence of predicted binding motifs alone does not demonstrate direct transcriptional regulation. Investigating E2F4 occupancy at these putative binding sites, as well as the downstream transcriptional networks, chromatin remodeling, and potential cofactors involved, would require extensive additional experimentation and is beyond the scope of the current work.

Our results demonstrate that TrkB signaling participates in the effects on proliferation, self-renewal and differentiation triggered by the E2F4DN-conditioned medium, as the blockage of TrkB signaling by TrkB-Fc resulted in the lack of effect of E2F4DN-conditioned medium on h5xFAD NSCs. Moreover, the inhibition of the TrkB receptor using ANA-12 also reflected the implication of the TrkB pathway in the proliferation improvement of the h5xFAD NSCs in presence of E2F4DN-conditioned medium, whereas no effect was observed in GFP-conditioned medium. Another piece of evidence came from the observation that E2F4DN-conditioned medium induced the upregulation of the transcripts encoding both TrkB (*Ntrk2*) and p75^NTR^ (*Ngfr*), likely as a readout of the presence of TrkB and p75^NTR.^-specific ligands in the medium (26). Finally, we demonstrated that TrkB is activated in the cerebral cortex of h5xFAD mice treated with E2F4DN, as evidenced by a significant increase in TrkB phosphorylated at Tyr516 in response to this treatment.

In contrast, the inhibition with THX-B of p75^NTR^, a receptor known to potentiate the differentiative effect of TrkB signaling on the oligodendrocyte lineage (26), could not significantly prevent the positive effect of the E2F4DN-conditioned medium on h5xFAD-derived NSC oligodendrogenesis. This result suggests that additional factors present in this conditioned medium may potentiate TrkB-dependent oligodendrogenesis in the absence of p75^NTR^ signaling. The effect of NT4/5 on the differentiation capacity of adult NSCs has not yet been fully characterized; therefore, the observed outcome in oligodendrocyte differentiation could reflect NT4/5’s specific action, which may differ functionally from BDNF despite both neurotrophins binding to and activating TrkB (80). Therefore, although our study identifies TrkB signaling as a key mediator of the neuronal E2F4DN-induced pro-neurogenic response, the precise secreted molecule(s) responsible for this activation require further investigation.

Several studies have evidenced a significantly decrease of TrkB early in AD (19). Our results have shown that both *Ntrk2* and *Ngfr,* are downregulated in h5xFAD SVZ and NSCs. Importantly, neuronal E2F4DN increased the expression of *Ntkr2* in h5xFAD NSCs and h5xFAD cortex, suggesting the promotion of TrkB signaling. Moreover, the E2F4DN treatment has proved to promote TrkB activation, evidenced by its specific phosphorylation. The implication of the TrkB pathway has been clearly demonstrated in multiple processes including neuronal survival, axonal outgrowth, cognitive function, or neurogenesis (26, 81), including in the AD context (82). A therapy targeting TrkB receptor upregulation, such as E2F4DN administration, could overcome the clinical limitations associated with BDNF as a therapeutic agent such as poor blood-brain barrier penetration or its short half-life (83). In fact, several studies have reported the beneficial role of TrkB in AD treatment (46, 84–85). Remarkably, unlike other therapies based on TrkB potentiation by agonists, E2F4DN therapy allows mitigating the AD phenotype through multifactorial targeting without affecting control individuals (83, 84). Although the mechanism implicated in TrkB up-regulation by E2F4DN in h5xFAD NSCs likely relies on the presence of NT4/5 in the conditioned medium since the activation of TrkB has been proved to induce upregulate the transcription of this receptor in WT NSCs (26), alternative mechanisms cannot be ruled out. Indeed, it has been shown that Aβ accumulation inhibits TrkB signaling (24, 86), and E2F4DN treatment decreases Aβ production and accumulation in 5xFAD mice (15); thus, the effect of neuronal E2F4DN in adult NSCs might be, at least partially, associated with indirect modulations such as Aβ reduction.

A key unresolved issue is the identity of the neuronal populations that influence the neurogenic niche. Evidence indicates that both cortical (87) and subcortical (73) neurons project to the SVZ and modulate adult neurogenesis. Notably, serotonergic axons arising from the raphe nucleus form close contacts with neural stem cells, promoting activity-dependent proliferation within the SVZ (73). Interestingly, these subcortical neurons, which are compromised in Alzheimer’s disease (88), are known to respond to NT4/5 signaling (89).

E2F4DN has shown to be a successful multifactorial agent for the different alterations occurring during AD pathogenesis (10, 12, 13, 15, 31). Importantly, this work provides a potential mechanism for these improvements, as previous studies have clearly demonstrated that the TrkB receptor is expressed across multiple brain cell types, highlighting TrkB signaling as a key pathway in the regulation of astrocytes, microglia, neurons and NSCs functions (26, 89–91).

### Limitations

The present study has some limitations. First, this study did not elucidate the molecular mechanism by which E2F4DN expression increases *Ntf5* expression in neurons. Defining this regulatory pathway was beyond the scope of the present work. Second, although we observed increased expression of the gene encoding NT4/5 in the absence of changes in BDNF levels—suggesting preferential activation of TrkB signaling via NT4/5—direct mechanistic evidence remains limited. Given the complexity of neurotrophin signaling and the potential involvement of additional TrkB ligands or parallel pathways, loss- and gain-of-function approaches will be necessary to establish the specific contribution of NT4/5 to TrkB activation in this context. Third, E2F4DN has previously been reported to reduce neuroinflammation and Aβ accumulation in the 5xFAD model (13, 15), processes that are closely linked to neurogenesis (1, 92). Therefore, although our findings support a role for TrkB signaling, it remains unclear whether TrkB activation represents the primary driver of the observed effects or one component of a broader, multifactorial mechanism underlying the therapeutic actions of E2F4DN.

### Conclusions

We have demonstrated that neuronal expression of a phosphorylation-resistant E2F4 mutant (Thr249/Thr251) restores ASN in the h5xFAD Alzheimer’s disease mouse model. This effect is associated with enhanced proliferation and differentiation of SVZ NSCs through activation of TrkB signaling (Fig. 7). Our findings identify neuronal E2F4 as a regulator of the neurogenic microenvironment, modulating paracrine signals that influence TrkB expression in NSCs. In this way, E2F4 coordinates communication between mature neurons and neural stem cells to fine-tune adult neurogenesis. These results support the therapeutic potential of targeting E2F4 phosphorylation to enhance neurogenesis in Alzheimer’s disease.

**Fig. 7.**
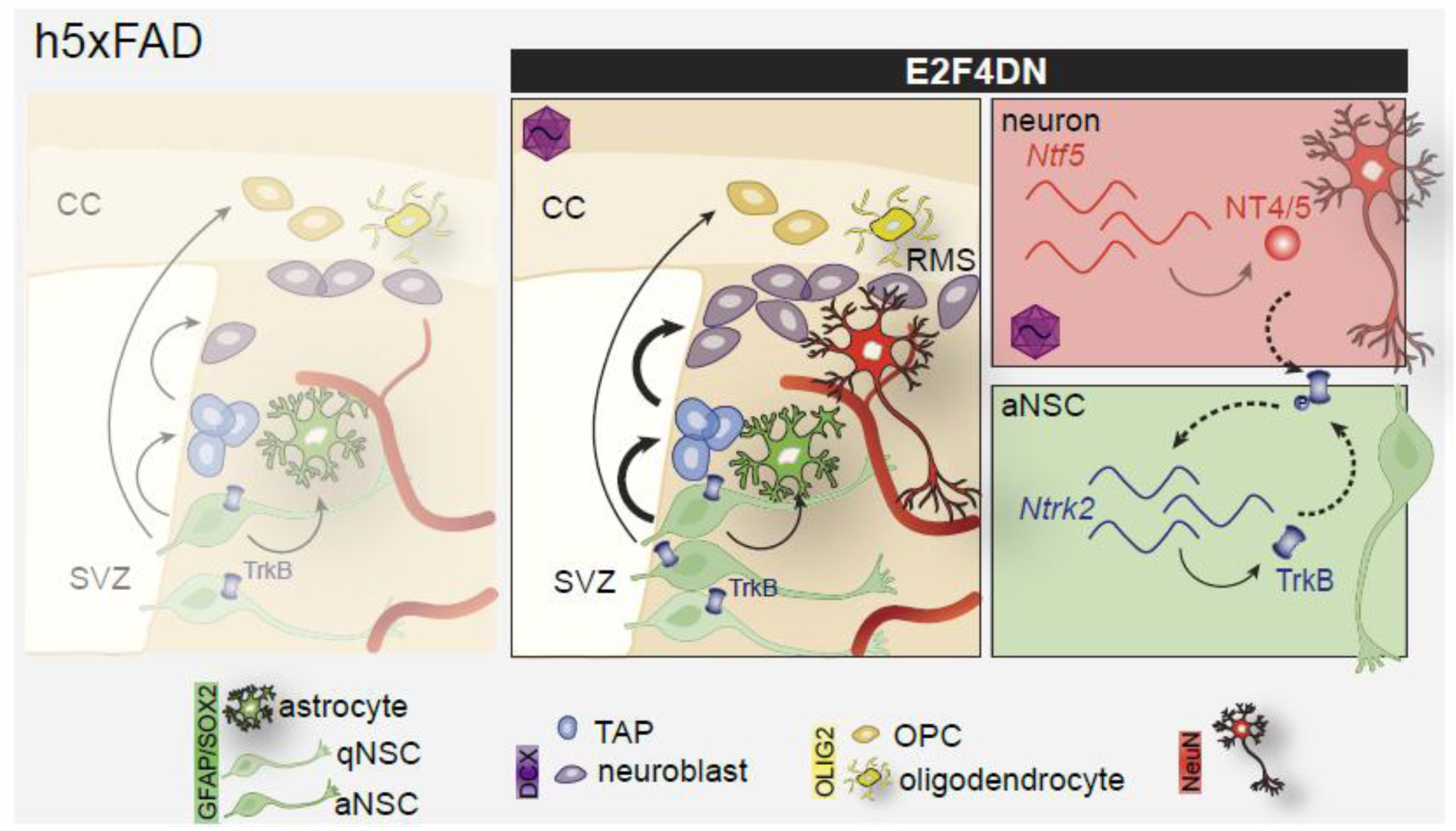
E2F4DN treatment restores the neurogenic populations in the SVZ of h5xFAD animals by promoting TrkB signaling. Adult neurogenesis is impaired in the SVZ of h5xFAD animals. The administration of AAV.E2F4DN in these animals increases the percentage of activated NSCs (aNSCs) in the SVZ, expanding the neuroblast population. Whereas the number of astrocytes in the parenchyma, and oligodendrocytes precursor cells (OPCs) and oligodendrocytes in the CC is unaltered after E2F4DN treatment. The neuronal expression of E2F4DN promotes the expression of the gene encoding TrkB in the adult SVZ-derived NSCs and the activation of TrkB pathway (P-TrkB). qNSC: quiescent neural stem cell; aNSC: activated neural stem cell; TAP: transcient amplifying precursor; OPC: oligodendrocyte precursor cell; CC: corpus callosum; SVZ: subventricular zone.

## Abbreviations

Aβ: amyloid β
AD: Alzheimer disease
Ala: alanine
ASN: adult subventricular neurogenesis
BDNF: brain-derived neurotrophic factor
FBS: fetal bovine serum
GFAP: glial fibrillary acidic protein
GLAST: glutamate transporter NSCs neural stem cells
NT4/5: neurotrophin 4/5
OB: olfactory bulb
PBS: phosphate buffer saline
PFA: paraformaldehyde
RMS: rostral migratory stream
RT: room temperature
SGZ: subgranular zone
Sox2: SRY-related HMG box 2
SVZ: subventricular zone
TAPs: transit-amplifying progenitors
Thr: threonine
TrkB: tropomyosin-related kinase
B WT: wild-type

## Declarations

### Ethics approval and consent to participate

All animal experiments were approved by the Animal Care and Ethics Committee of the Consejo Superior de Investigaciones Científicas (CSIC) and the Autonomous Government of Madrid and were conducted in accordance with Spanish and European Union regulations on animal experimentation.

Informed consent was not applicable to this study.

### Consent for publication

Not applicable.

### Availability of data and materials

All data generated or analyzed during this study are included in this published article and its supplementary information files.

### Authors’ contributions

Design of the work: ALU and JMF. Acquisition of data: ALU, AMLM, MRL, AF, CSP, AGG, VCD, and LVL. Analysis and interpretation of data: ALU and JMF. The manuscript was written by ALU and JMF. All authors commented on previous versions of the manuscript. All authors read and approved the final manuscript.

## Acknowledgements

We acknowledge the technical support provided by the Microscopy and Animal Facility of the Cajal Institute.

## Authors’ information

Current address for ALU: Tetraneuron.

