## Supplementary Material for "Neuronal expression of E2F4DN restores adult neurogenesis in homozygous 5xFAD mice via TrkB signaling"

Corresponding author: José María Frade

**This PDF file includes:**

Figures S1 to S6

Tables S1 to S3

**Figures S1 to S6**

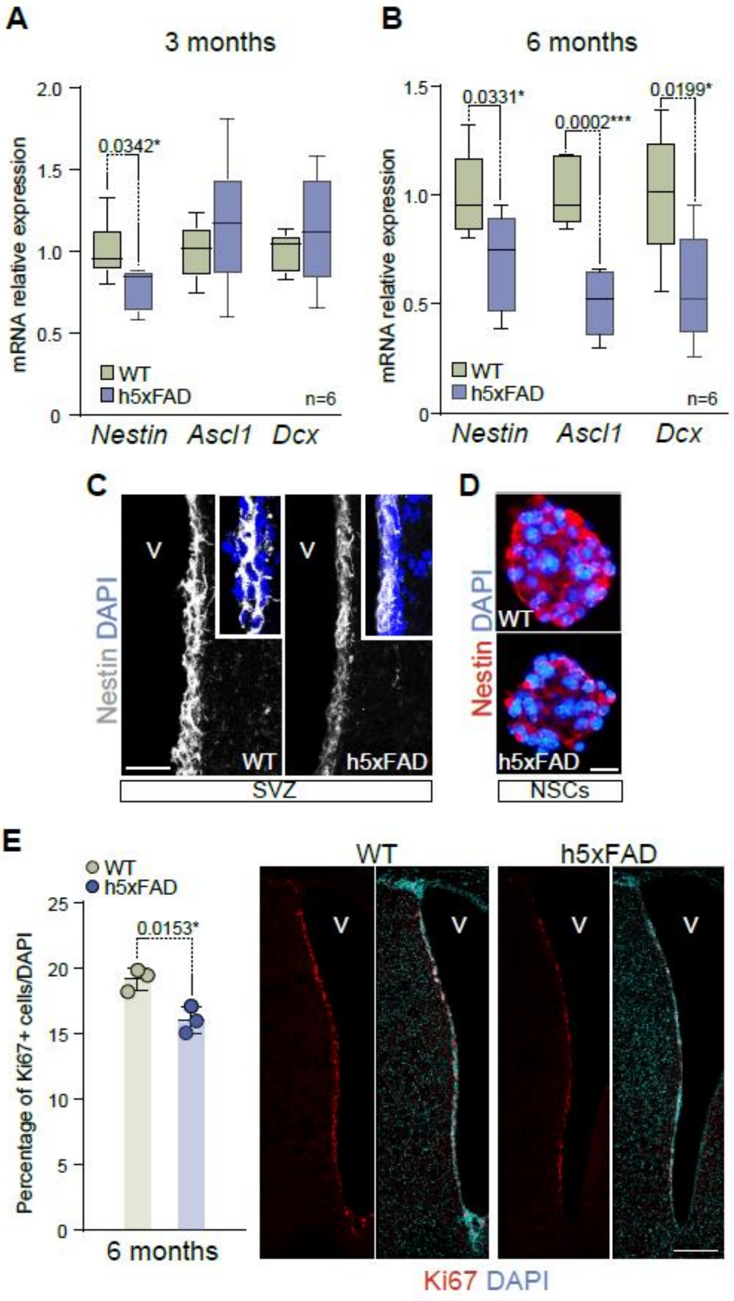


**Fig. S1** Neurogenic alterations in the SVZ of h5xFAD mice worsen with age. **A** Gene expression of *Nestin*, *Ascl1* and *Dcx* in the SVZ of 3-month-old WT (green) and h5xFAD (blue) mice. *Rpl27* was used for normalization (two-tailed Student's t-test). **B** Gene expression of *Nestin*, *Ascl1* and *Dcx* in the SVZ of 6-month-old WT (green) and h5xFAD (blue) mice. *Rpl27* was used to normalize data (two-tailed Student's t-test). **C** Representative images of the immunohistochemistry for Nestin in the SVZ of WT and h5xFAD animals. **D** Representative images of the immunohistochemistry for Nestin in the NSCs cultures of WT and h5xFAD animals. **E** Percentage of Ki67 positive cells in the SVZ of 6-month-old WT and h5xFAD animals (left panel) (two-tailed Student's t-test). Representative images of the immunohistochemistry for Ki67 (right panel). DAPI was used to counterstain DNA. V: ventricle. Scale bar: 25 μm (C); 10 μm (D); 200 μm (E).

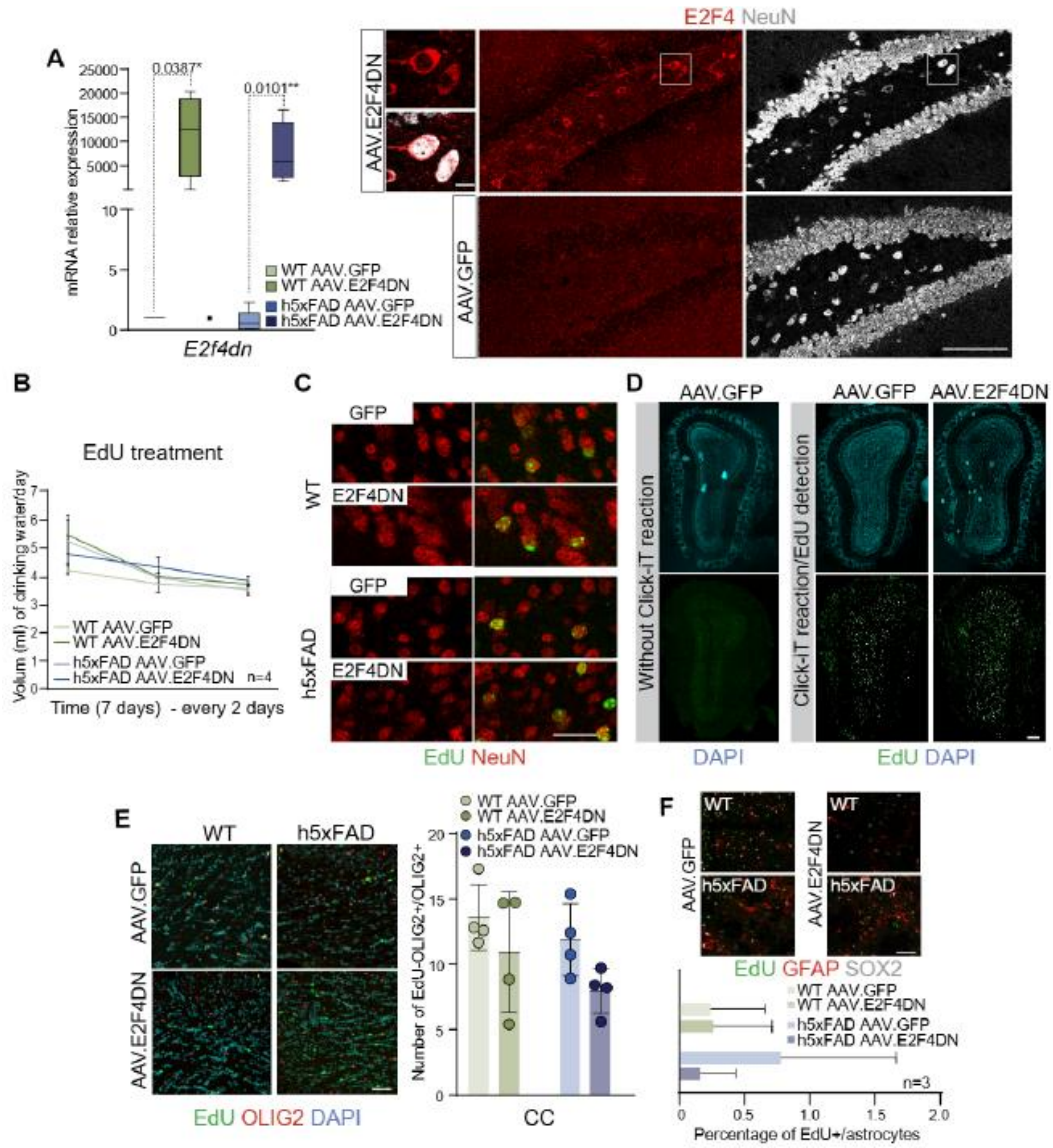


**Fig. S2** EdU treatment to analyze NSCs behavior *in vivo*. **A** Expression of the E2F4DN construct in the SVZ of WT and h5xFAD mice 1 month after tail vein injection with AAV.GFP or AAV.E2F4DN. *Rpl27* was used for normalization (left panel) (one-way ANOVA followed by Tukey's post-hoc test and). Representative images of the immunohistochemistry for E2F4 and NeuN in E2F4DN and GFP-treated animals (right panel).

**B** Recording of drunk water during the 7 days of EdU treatment in WT and h5xFAD animals injected with GFP or E2F4DN (one-way ANOVA followed by Tukey's post-hoc test and). **C** Immunohistochemistry for NeuN and EdU in the OB of WT and h5xFAD animals 1 month after injection with AAV.GFP (GFP) or AAV.E2F4DN (E2F4DN). **D** EdU detection in the OB of WT mice injected with AAV.GFP or AAV.E2F4DN (left panel). OB section of a photobleached WT mouse (right panel) previous to EdU detection with Click-iT reaction (right panel). **E** Number of OLIG2+ cells with EdU labelling in the corpus callosum (CC) of WT and h5xFAD mice injected with AAV.GFP or AAV.E2F4DN (right panel) (one-way ANOVA followed by Tukey's post-hoc test and). Left panel: representative images. **F** Immunohistochemistry for GFAP and SOX2 astrocytes with EdU labellig in the striatum of WT and h5xFAD animals treated with AAV.GFP or AAV.E2F4DN. The percentage of EdU+/GFAP+/SOX2+ cells are indicated in the images (Kruskal-Wallis non-parametric test). DAPI was used to counterstain DNA. Scale bars: 100 μm, and 10 μm in the amplification, (A); 50 μm (C, E, F); 200 μm (D).

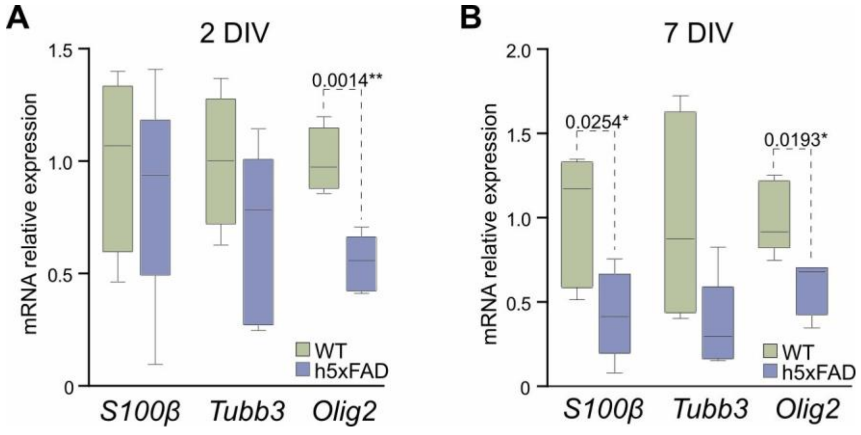


**Fig. S3** Differentiation is impaired in h5xFAD NSCs. **A** Gene expression of *S100β, Tubb3* and *Olig2* in WT and h5xFAD NSCs after 2 DIV under differentiation conditions (two-tailed Student's t-test). **B** Gene expression of *S100β, Tubb3* and *Olig2* in WT and h5xFAD NSCs after 7 DIV under differentiation conditions (two-tailed Student's t-test). *Rpl27* was used to normalize data.

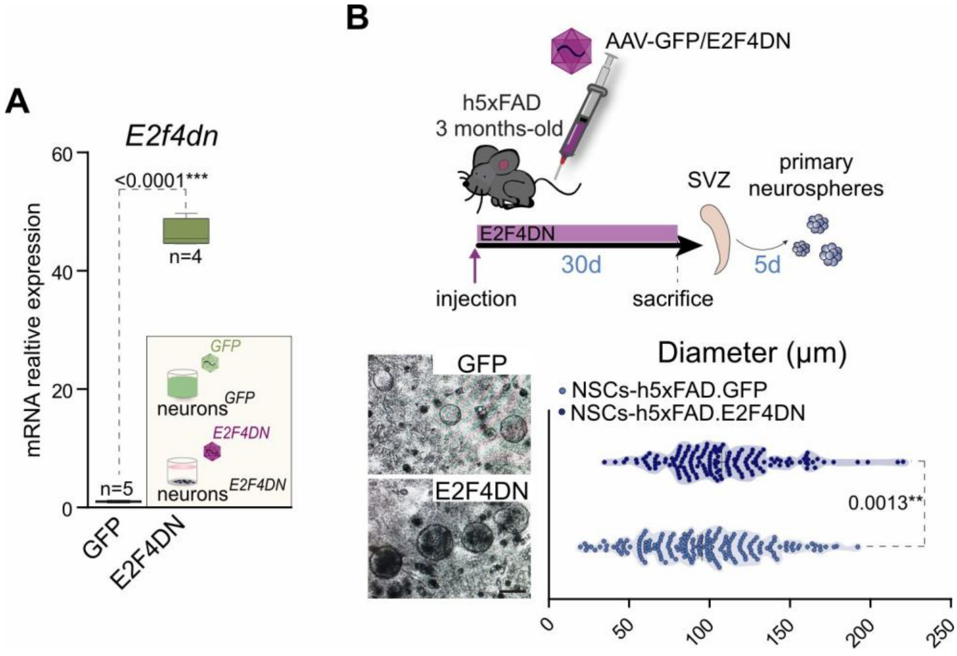


**Fig. S4** Diameter of neurospheres increases in h5xFAD mice injected with AAV.E2F4DN. **A** *E2f4dn* gene expression in cortical cultures 48h after transduction with Ad5.GFP (GFP) or Ad5.E2F4DN (E2F4DN) (two-tailed Student's t-test). *Rpl27* was used for normalization. **B** Scheme of the generation of primary neurospheres from the SVZ of GFP- or E2F4DN-injected mice. Three-month old animals were injected with GFP or E2F4DN and, 1 month later, SVZs were isolated and NSCs were cultured during 5 DIV under proliferating conditions to analyze their behavior (upper panel). Diameter of the neurospheres isolated from GFP- or E2F4DN-injected h5xFAD animals (bottom panel) (two-tailed Student's t-test). Representative images are shown. Scale bars: 100 μm (B).

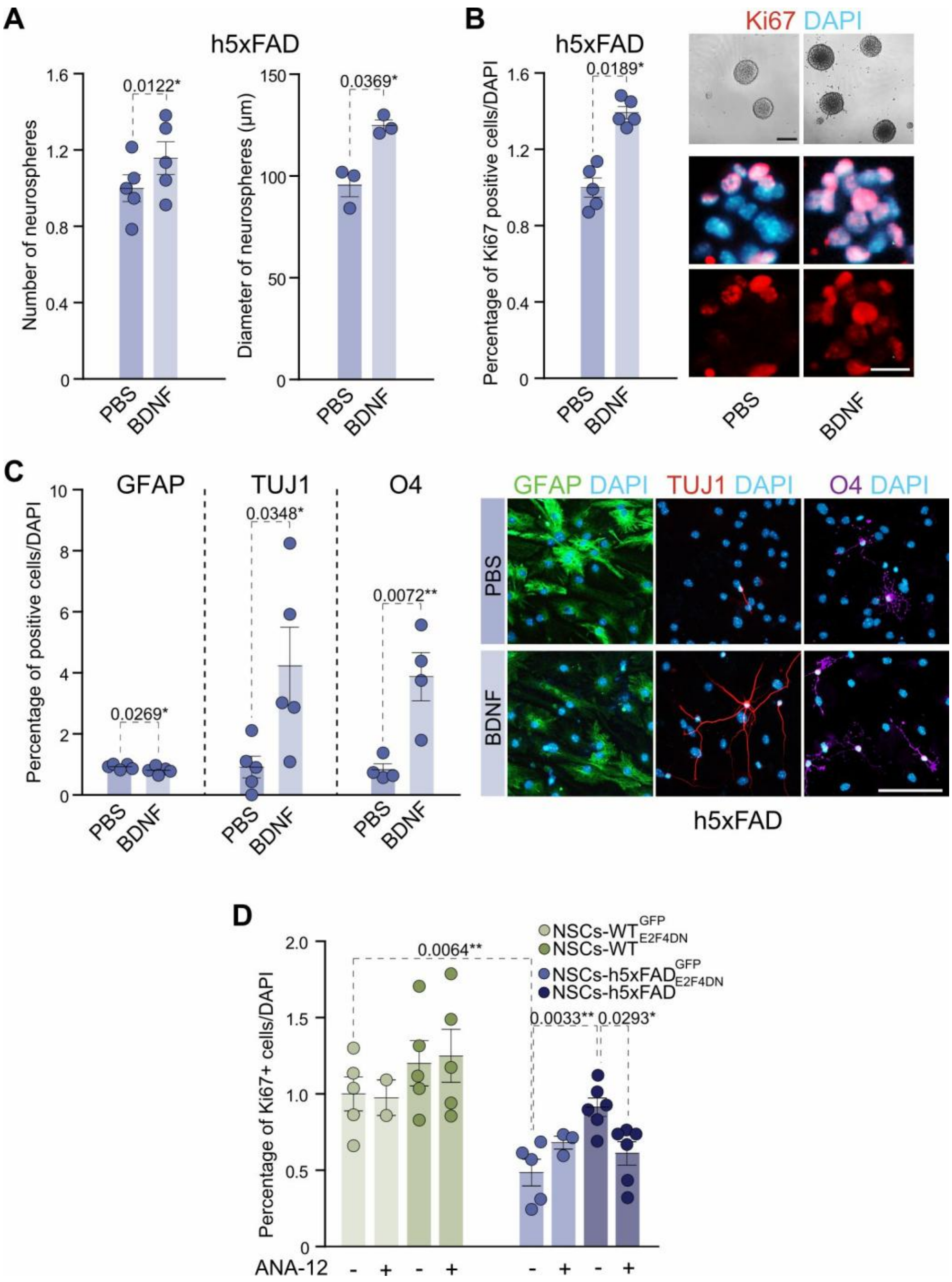


**Fig. S5** BDNF promotes self-renewal, proliferation and differentiation of h5xFAD NSCs. **A** Number (left panel) and diameter (right panel) of neurospheres in untreated (PBS) and BDNF treated h5xFAD NSCs (two-tailed Student's t-test). **B** Percentage of Ki67 positive cells in h5xFAD NSCs treated or untreated with BDNF (left panel) (two-tailed Student's t-test). Representative images are shown (right panel). **C** Percentage of astrocytes (GFAP), neurons (TUJ1) and oligodendrocytes (O4) in control (PBS) or BDNF-treated h5xFAD NSCs after 7 DIV under differentiation conditions (left panel) (two-tailed Student's t-test). Representative images are shown (right panel). DAPI was used to counterstain DNA. **D** Percentage of Ki67+ WT NSCs or h5xFAD NSCs grown for 5DIV with GFP- or E2F4DN-conditioned medium in presence of the TrkB antagonist ANA-12 (one-way ANOVA followed by Tukey's post-hoc test and). DAPI was used to estimate the total number of cells. Scale bars:100 µm (B; bright-field); 10 µm (B; immunofluorescence); 50 µm (C).

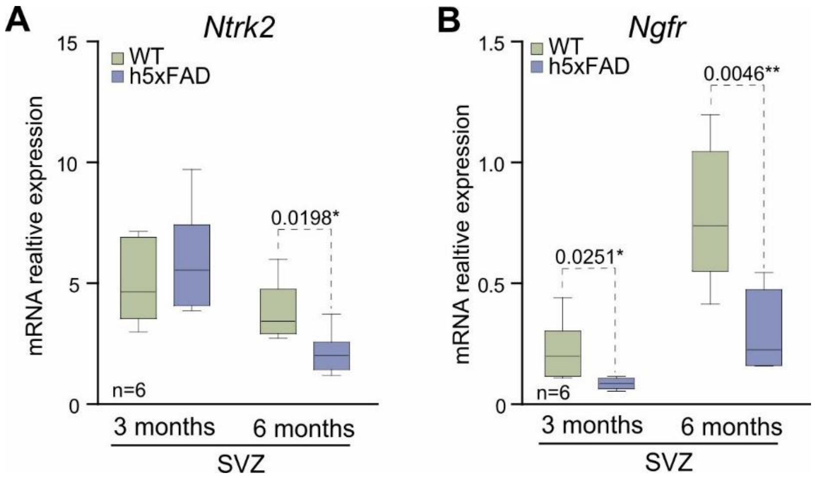


**Fig. S6** *Ntrk2* and *Ngfr* expression in the SVZ of h5xFAD mice. **A** qPCR expression of *Ntrk2* in the SVZ of h5xFAD and WT mice of 3 months (left panel) and 6 months (right panel) of age (two-tailed Student's t-test). **B** qPCR expression of *Ngfr* in the SVZ of h5xFAD and WT mice of 3 months (left panel) and 6 months (right panel) of age (two-tailed Student's t-test). *Rpl27* was used for normalization in both cases.

### Tables S1 to S3

**Table S1. List of primary antibodies for immunocytochemistry (ICC), immunohistochemistry (IHC) and western blot (WB).**

| Protein | Host | Application | Dilution | Source | Reference |
|---|---|---|---|---|---|
| **Actin** | Mouse | WB | 1:10,000 | Millipore | MAB1501R |
| **BDNF** | Mouse | ICC | 1:200 | Icosagen | 327-100 |
| | Rabbit | ICC; IHC; WB | 1:200;1:200;1:1,000 | Proteintech | 28205-1-AP |
| **Dcx** | Rabbit | IHC | 1:400 | Synaptic Systems | 326 003 |
| **E2F4** | Mouse | IHC | 1:100 | Millipore | MABE160 |
| **GFAP** | Rabbit | ICC; IHC | 1:800 | Abcam | AB7260 |
| **Ki67** | Rabbit | ICC; IHC | 1:800;1:600 | Abcam | AB16667 |
| | Rat | IHC | 1:600 | | |
| **MAP2** | Chicken | ICC | 1:500 | Abcam | AB5392 |
| **Nestin** | Mouse | ICC | 1:600 | Abcam | AB6142 |
| **NeuN** | Rabbit | IHC | 1:400 | Millipore | MAB377 |
| **O4** | Mouse | ICC | 1:600 | R&D Systems | MAB1326 |
| **Olig2** | Rabbit | IHC | 1:100 | Merk | AB9610 |
| **Sox2** | Goat | IHC | 1:300 | R&D Systems | AF2018 |
| **TrkB** | Mouse | WB | 1:1,000 | BD Biosciences | 610102 |
| **P-TrkB (Tyr516)** | Rabbit | WB | 1:1,000 | Invitrogen | PA5-36695 |
| **TUJ1** | Mouse | ICC | 1:300 | Millipore | MAB5564 |

**Table S2. List of secondary antibodies for ICC, IHC and WB.**

| Antibody | Host | Application | Dilution | Source | Reference |
|---|---|---|---|---|---|
| **488 anti-rabbit** | Donkey | ICC; IHC | 1:800 | Invitrogen | A21206 |
| **555 anti-rat** | Donkey | IHC | 1:800 | Invitrogen | 17236543 |
| **568 anti-rabbit** | Donkey | ICC; IHC | 1:800 | Invitrogen | 10617183 |
| **594 anti-mouse** | Goat | ICC; IHC | 1:800 | Invitrogen | A11032 |
| **647 anti-chicken** | Goat | ICC | 1:800 | Invitrogen | A21449 |
| **647 anti-goat** | Donkey | IHC | 1:800 | Jackson | A32849 |
| **647 anti-mouse** | Donkey | ICC; IHC | 1:800 | Invitrogen | A31571 |
| **680RD anti-mouse** | Goat | WB | 1:14000 | LI-COR | 926-68070 |
| **800CW anti-rabbit** | Goat | WB | 1:14000 | LI-COR | 925-32211 |

**Table S3. List of primers for qPCR analysis.**

| Gene | | Sequence (5'→3') |
|---|---|---|
| *Ascl1* | Fw | GACTTTGGAAGCAGGATGGCA |
| | Rv | ACCCCTGTTTGCTGAGAACAT |
| *Bdnf* | Fw | ATTACCTGGATGCCGCAAAC |
| | Rv | CAGTTGGCCTTTGGATACCG |
| *Dcx* | Fw | ACACCCTTGATGGAAAGCAG |
| | Rv | TGTTCATTGCTTGTGGTCCT |
| *E2f4dn (AAV)* | Fw | CCCGGAGACCACGATTACAT |
| | Rv | AAGCAGCGTATCCACATAGC |
| *E2f4dn (Ad5)* | Fw | AGCCATTCCCTGGACTTCT |
| | Rv | TCCTCAGCTCGCTCCCG |
| *Nestin* | Fw | CTGCAGGCCACTGAAAAGTT |
| | Rv | GACCCTGCTTCTCCTGCTC |
| *Ntf5* | Fw | CAAGGCTAAGCAGTCCTATGT |
| | Rv | CAGTCATAAGGCACGGTAGAG |
| *Ntrk2 (TrkB)* | Fw | CAGTATTAACTCGCTTCTGGC |
| | Rv | TTCATCCACGTCAAAGGCAG |
| *Olig2* | Fw | CGCAAGCTCTCCAAGATCG |
| | Rv | CTCACCAGTCGCTTCATCTC |
| *Ngfr ($p75^{NTR}$)* | Fw | CTAGGGGTGTCCTTTGGAGGT |
| | Rv | CAGGGTTCACACACGGTCT |
| *Rpl27* | Fw | CCCTCCTTTCCTTTCTGCTG |
| | Rv | GCCATCGTCAATGTTCTTCAC |
| *S100β* | Fw | AAAGTGATGGAGACGCTGGA |
| | Rv | CTTTGCTGTGCCTCCTCTTG |
| *Tubb3* | Fw | TCCGAGTACCAGCAGTACCA |
| | Rv | GGCTTCCGATTCCTCGTCAT |
